# Lactate potentiates NMDA receptor currents via an intracellular redox mechanism targeting cysteines in the C-terminal domain of GluN2B subunits: implications for synaptic plasticity

**DOI:** 10.1101/2024.11.21.624499

**Authors:** Hubert Fiumelli, Gabriel Herrera-López, Fouad Lemtiri-Chlieh, Lorène Mottier, John Girgis, Carine Ben-Adiba, Pascal Jourdain, Nicolò Carrano, Hanan Mahmood, Amanda Ooi, Stefan T. Arold, Monica Di Luca, Fabrizio Gardoni, Pierre J. Magistretti

## Abstract

Through the Astrocyte Neuron Lactate Shuttle, astrocyte-derived lactate fuels the high-energy demands of neurons and acts as a signaling molecule, promoting synaptic plasticity and memory consolidation. Lactate regulates neuronal excitability and modulates the expression of genes related to synaptic plasticity and neuroprotection, but the molecular mode for these signaling actions is uncertain. Using patch-clamp recordings in cultured cortical neurons, we found that lactate enhances both the amplitude and the inactivation time constant of NMDA receptor currents (I_NMDAR_) evoked by brief applications of glutamate and glycine. Not reproduced by HCAR1 agonists, this modulation depends on monocarboxylate transporters and lactate dehydrogenase, indicating the requirement for lactate entry and metabolic conversion into pyruvate and NADH formation within neurons. Disruption of intracellular calcium dynamics or inhibition of Ca^2+^/calmodulin-dependent protein kinase II (CaMKII), a NMDAR-associated kinase linking Ca^2+^ signal to long-term potentiation (LTP), significantly diminishes the effects of lactate on I_NMDAR_. We identified two redox-sensitive cysteine- containing sequences in the intrinsically disordered intracellular C-terminal domain of the GluN2B subunit that play a role in the potentiation of NMDAR by lactate. In a compelling set of experiments using HEK cells, we observed that the presence of functional CaMKII and GluN2B-containing NMDARs is necessary for the lactate-enhancing effects. Mutations in GluN2B that prevent CaMKII binding or redox regulation via cysteines abrogate the modulatory action of lactate. Immunoprecipitation experiments in neurons attest that lactate increases the association between CaMKII and GluN2B. This interaction is crucial for the potentiation of I_NMDAR_ amplitude by lactate. Proximity ligation assays between GluN2B and the postsynaptic density marker PSD-95 revealed that lactate induced an accumulation of GluN2B in dendritic spines, an effect that was prevented by a CaMKII peptide inhibitor. These results highlight a mechanistic pathway whereby lactate boosts NMDAR function through intracellular metabolic conversion and redox-sensitive interactions requiring CaMKII, establishing a link between astrocyte metabolism and synaptic modulation in neurons.

## Introduction

Lactate is not only a preferred energy substrate to sustain neuronal activity, but it is also emerging as an important signaling molecule regulating higher brain functions (Machler et al., 2016; Zimmer et al., 2017; Magistretti and Allaman, 2018; Zuend et al., 2020; Roumes et al., 2021; Dembitskaya et al., 2022). Lactate signaling plays a role in learning and memory (Suzuki et al., 2011; Akter et al., 2023), sleep (Petit et al., 2015), mood disorders (Carrard et al., 2021), and stress adaptation (Karnib et al., 2019). In response to increased neuronal activity engaged in higher brain functions such as learning and memory (Dembitskaya et al., 2022), glial cells upregulate their glucose uptake and aerobic glycolysis, resulting in the production of lactate that is passed on to neurons via monocarboxylate transporters (MCTs) in a process known as the astrocyte-to-neuron lactate shuttle (ANLS) (Pellerin and Magistretti, 1994; Magistretti and Allaman, 2015). Subsequently, in neurons, lactate is oxidized to pyruvate by the action of lactate dehydrogenase-1, concomitantly reducing NAD^+^ to NADH. In mitochondria pyruvate feeds the tricarboxylic acid cycle and oxidative phosphorylation to produce adenosine triphosphate (ATP). Along its metabolic journey to support neurons, astrocyte-derived lactate can also elicit many non-metabolic effects to support the function of neurons and other cells (Magistretti and Allaman, 2018). For instance, it has been shown to protect neurons from oxidative stress through hormetic reactive oxygen species (ROS) signaling (Tauffenberger et al., 2019) or from excitotoxicity resulting from excessive glutamate exposure through a coordinated cellular pathway involving ATP production, release, and activation of a P2Y receptor/K_ATP_ cascade (Jourdain et al., 2016; Jourdain et al., 2018). A growing number of studies performed in several species have convincingly demonstrated that lactate originating from the coupling between astrocytes and neurons is necessary for learning and memory (Newman et al., 2011; Suzuki et al., 2011; Harris et al., 2019; Akter et al., 2023; Frame et al., 2023). Blocking the production of lactate from astrocytes and/or its delivery to neurons, and supplying exogenous lactate, not only hampers, and respectively rescues, memory formation (Suzuki et al., 2011; Boury-Jamot et al., 2015; Vezzoli et al., 2020) but also affect the electrophysiological and morphological correlates of learning such as changes in long-term potentiation (LTP) and spine density. In this context, lactate contributes to the excitability (Karagiannis et al., 2021) of neuronal networks and supports long-term potentiation in the rodent hippocampus (Suzuki et al., 2011). The contribution of lactate in long-term memory formation relies on modulation of synaptic plasticity and engages activity-dependent genes (Suzuki et al., 2011). Yang et al. (2014) found that lactate induces the expression of well-known immediate-early genes related to plasticity, such as Arc, Zif268 and c-Fos, through a mechanism involving N-methyl-D- aspartate-type glutamate receptors (NMDAR), calcium and the MAPK signaling pathway. Interestingly, a large proportion of genes regulated by lactate depends on NMDAR activity and are similarly modulated by NMDA and NADH (Margineanu et al., 2018), but not by the equicaloric glycolytic co-end product pyruvate. These observations suggest that changes in the redox state, possibly due to an increase in the NADH/NAD^+^ ratio upon lactate uptake and conversion to pyruvate by the lactate dehydrogenase, may mediate the effect of lactate on NMDARs. Ca2+/calmodulin-dependent protein kinase II (CaMKII), the prototypical mediator of cellular Ca2+ signals and central regulator of neuronal plasticity, interacts with NMDARs at synapses (Gardoni et al., 1998; Halt et al., 2012) and is sensitive to the redox state (Erickson et al., 2008; Bayer and Schulman, 2019) as is the NMDAR (Choi and Lipton, 2000; Bodhinathan et al., 2010). Building on the foregoing experimental findings, we have undertaken a series of studies to unravel the molecular mechanism through which lactate modulates NMDARs and tested whether CaMKII participates in the molecular pathway mediating the effects of lactate on neuronal plasticity.

## Material and Methods

### DNA constructs and point mutants

Our experimental approach involved the use of pcDNA3.1 plasmids harboring the mouse cDNA for Grin1 (OMu21895, accession NM_008169), Grin2a (OMu22286, accession NM_008170) or Grin2b (OMu22276, accession NM_008171.3). These plasmids were sourced from GenScript. In addition, we obtained a pCMV6-Entry vector expressing CaMKIIα (MR207662, accession NM_177407) from OriGene. The pcDNA3.1-Peredox mCherry construct was a gift from Gary Yellen (Addgene #32383).

Multiple point mutants of the Grin2b and CaMKII constructs were generated by GenScript. Specifically, we engineered two pcDNA3.1-Grin2b double mutants to prevent CaMKII binding, based on previous studies (Strack et al., 2000; Halt et al., 2012). The first double mutant includes L1298A and R1300Q substitutions, while the second double mutant contains R1300Q and S1303D substitutions. Furthermore, the pcDNA3.1-Grin2b construct was modified to produce two redox mutants bearing multiple cysteine substitutions: redox- sensitive site 1 mutant with C946S and C954S, and redox-sensitive site 2 mutant with C1215S, C1218S, C1239S, C1242S, and C1245S. Lastly, we designed a vector expressing a redox-insensitive (Δredox) CaMKII triple mutant incorporating M281S, C280S, and C289S substitutions. These amino acid switches were selected based on previous findings on the redox modulation of CaMKII activity (Erickson et al., 2008; Coultrap et al., 2014; Rocco- Machado et al., 2022).

### Cell culture and generation of stable cell lines

Human embryonic kidney 293T/17 (HEK) cells were obtained from American Type Culture Collection (CRL11268) and cultured in DMEM (ThermoFisher, 31966) supplemented with 10% fetal bovine serum, 100 U/mL penicillin, and 100 mg/mL streptomycin in a humidified atmosphere of 95% air and 5% CO2 at 37°C. Cells were subcultured when they reached a confluence of around 80-90%. Under normal conditions, HEK cells do not express NMDAR subunits or CaMKIIα (figure S2D) and represent a good pharmacogenetic model to study ions channel functions (Kohr and Seeburg, 1996; Thomas and Smart, 2005; Ooi et al., 2016; Zhang et al., 2022). To express functional NMDARs, cells were grown to approximately 50% confluence on poly-D-lysine-coated glass coverslips (Neuvitro) and transfected using 1 µg of a cocktail of expression vectors encoding GluN1, GluN2a or GluN2b, and GFP, mixed at a 1:1:0.25 ratio, and 2 µl of Lipofectamine 2000 (Invitrogen, 11668-019) according to the manufacturer’s instructions. GFP serves as a marker to aid finding cells likely to express NMDARs when visualized with fluorescence microscopy. Ketamine (500 µM), or a mix of D- AP5 (50 µM, abcam) and 7-chloro kynurenic acid (200 µM, abcam) was added to the culture media to prevent excitotoxicity due to the functional expression of NMDAR. To generate stable CaMKIIα-expressing cell lines, HEK cells (1 x 10^6^) were seeded in 60 mm cell culture dishes and transfected 24h later at a confluence of 50-70% with 6 µg of DraIII-linearized pCMV6-Entry wild-type or Δredox mutant CaMKIIα expression vector using lipofectamine 3000 (Invitrogen) according to the manufacturer’s instructions. Clones were selected from cultures grown for 14 days in the presence of 800 µg/ml G418 sulfate (Invitrogen).

### Cultured cortical neurons

Primary cultures of cortical neurons were prepared from CD-1 IGS mouse embryos at embryonic day 17 (sourced from Charles River Laboratories, UK). This process followed a previously described protocol (Allaman et al., 2010), with minor modifications. Procedures were conducted in accordance with the Guide for the Care and Use of Laboratory Animals (8^th^ edition, National Research Council, 2011) and Implementing Regulations of the Law of Ethics of Research on Living Creatures (3^rd^ ed, National Committee of BioEthics, 2022). They were performed under the oversight of the local Institutional Animal Care and Use Committee under protocol number: 23IACUC007). Briefly, minced pieces (1-2 mm^3^) of isolated cerebral cortices were enzymatically digested in Hank’s buffered saline solution containing papain (20 U/ml, Worthington Biochemical), L-cysteine (1 mM, Sigma-Aldrich), DNase I (100 µg/ml, Worthington Biochemical) and EDTA (0.5 mM, Sigma-Aldrich), and gently triturated to a single-cell suspension. Dissociated cells, filtered over a 40 μm cell strainer (Falcon), were plated onto poly-L-ornithine-coated cell culture dishes at a density of 4 x 10^4^ cells/cm^2^ and maintained in Neurobasal medium supplemented with 1x B27, L-glutamine (0.5 mM), penicillin (50 U/mL), and streptomycin (50 U/mL) (all from Gibco) at 37 °C in a humidified atmosphere of 5% CO_2_ and 95% air. Neurons were used for experiments at day in vitro (DIV) 11-17, except for co-immunoprecipitation and proximity ligation assays, which utilized neurons at DIV 14-17.

### Patch Clamp recordings from cultured neurons and HEK cells

Patch pipettes were pulled from thin-walled borosilicate glass capillaries with filament (BF150- 110-10HP, Sutter Instrument®, Novato, CA) using a P-1000 Flaming/Brown micropipette puller (Sutter Instrument®, Novato, CA). Pipette resistances were between 3 to 6 MΩ when filled with an intracellular solution consisting (in mM) of 130 CsCl, 1 CaCl2, 1 MgCl2, 10 EGTA, 10 HEPES, 2 Na-ATP, 0.3 Na-GTP, and 10 phosphocreatine (pH 7.30 adjusted with Trizma® base or CsOH, and osmolarity 290 ± 5 mOsm adjusted with sucrose). Cells were perfused at a rate of 1-2 ml/min with an external bath solution consisting of (in mM): 125 NaCl, 2.5 KCl, 2 CaCl_2_, 1 MgCl_2_, 10 HEPES, 10 glucose (pH 7.30 adjusted with Trizma® base or NaOH, osmolality 300 ± 5 mOsm adjusted with sucrose). The extracellular solution was supplemented with reagents at concentrations indicated in the text, which was matched with sodium-gluconate or NaCl in the control to offset the change in osmolarity. Upon achieving whole-cell configuration, the cells were voltage-clamped at a holding potential of - 30 mV. Data were acquired using a MultiClamp 700B amplifier (Molecular Devices). Currents were low pass filtered at 2 kHz before analog-to-digital conversion, digitized at 20 kHz, and uncorrected for leakage current. Data acquisition and analysis were achieved with pClamp (version 10.7). Series resistance and cell capacitance were compensated and monitored throughout the recordings. Experiments were discarded if the series resistance changed by more than 20%. NMDAR currents were evoked using brief pulses of external solution containing glutamate (50 μM) and glycine (100 μM) from a micropipette positioned approximately 50 µm away from the recorded cell. Stimulations were controlled by air puffs (1 ms, 15 psi) delivered using a Picospritzer III (Parker Hannifin, Hollis, NH). To isolate NMDAR responses in neuronal cultures, we blocked non-NMDA ionotropic glutamate receptors with 6- cyano-7-nitroquinoxaline-2,3-dione (CNQX, 50 µM). We also blocked voltage-dependent sodium channels with tetrodotoxin (TTX, 0.5 µM) to avoid unwanted NMDAR responses triggered by spontaneous action potentials. Current (or charge) to voltage relationships (I-V or Q-V) for NMDARs were constructed from recordings using a protocol consisting of a series of 15 s voltage depolarization between +60 to -40 mV in 20 mV decrements. NMDAR stimulations were triggered 4 s after the beginning of the voltage clamp protocol. Tau (*τ*) values for the inactivation phase of NMDAR currents (I_NMDAR_) were approximated by fitting the signals (10% - 90%) with a single exponential decay equation I(t) = α * e^-t/^*^τ^* (using the Clampfit module). Cultured neurons were recorded at DIV 11-17 and recordings from HEK cells were performed 18 to 30hr after the transfection

### NADH/NAD+ measurements

HEK cells were plated on poly-D-lysine-coated glass coverslips (Neuvitro), and transfected with a plasmid encoding peredox-mCherry (Hung et al., 2011; Hung and Yellen, 2014). One to two days later, cells were first equilibrated for 15-30 minutes and then imaged in a solution consisting of (in mM) 140 NaCl, 3 mM KCl, 2 CaCl2, 1 MgCl2, 5 glucose, and 10 HEPES, pH 7.35. Epi-fluorescence images were acquired with a Zyla 5.5 sCMOS camera (Andor) at a frame rate of 0.1Hz on a DHM T1000 microscope equipped with a custom fluorescence module (Pavillon et al., 2010) controlled with micro-Manager (Edelstein et al., 2010). Successive illuminations of 60-100ms at 395 and 574 nm from a monochromator (Polychrome V, Till Photonics) with a 15 nm slit were used to excite Peredox and mCherry, respectively. Green or red fluorescence was collected through a triple-edge dichroic beamsplitter and an emission filter (524/24 nm or 628/32 nm, Semrock) mounted on a motorized wheel (Prior Scientific).

Image analysis was performed in FIJI. We first corrected the timelapse x-y drift, if any, with the Correct 3D Drift plugin (Parslow et al., 2014), subtracted the background, and generated a Peredox to mCherry pixel-by-pixel ratio at each time point. Region of interests over whole cells expressing the sensor were delineated with a custom segmentation macro, and the average Peredox/mCherry ratio for each cell was normalized to that of the baseline period.

### Calcium imaging

Calcium imaging experiments were performed as described by Jourdain et al. (2018). In brief, neurons were loaded with 4 µM Fura-2 AM (Molecular Probes) for 30 min at 37°C in HBSS (consisting of in mM: 140 NaCl, 3 KCl, 2 CaCl2, 1 MgCl2, 5 glucose, 10 HEPES, adjusted to pH 7.3) containing 1% bovine serum albumin (Sigma) in the presence or not of CaMKII inhibitors. Coverslips were then mounted in a 1 ml closed chamber and perfused at a rate of 2 ml/min with HBSS. Imaging was performed using a DHM T1000 inverted microscope (Lyncée Tec, Switzerland) equipped with a custom fluorescence module. The ratiometric dye Fura-2 was successively excited at 340 and 380 nm using a monochromator (Polychrome V, Till Photonics), and the emission light was collected at 510 nm (at bandwidth of 84 nm) by a CCD camera (INFINITY 3S-1, Lumenera Corporation). Image pairs (1392 x 1040 pixels) were acquired every 6.5 sec (150 ms exposures, 2 x 2 binning) with the micro-Manager software (Edelstein et al., 2010), and the background-subtracted fluorescence images were further processed with ImageJ. The baseline-subtracted averaged fluorescence intensity ratio of images excited 340 over 380 nm from a 50-pixel selection centered over neuron soma was used to monitor changes in intracellular Ca^2+^ concentration in response to 1-2 min pulses of 1 μM glutamate and 100 μM glycine.

### Intracellular pH measurements

For these experiments, cells were seeded at a density of 10,000 cells per well in 96-well plates (Corning, 3340), reaching approximately 90% confluency 48h later when they were incubated for 30 min at 37°C with the dual excitation ratiometric pH indicator BCECF-AM (2.5 μM, Invitrogen) in HBSS, pH 7.3. Subsequently, cells were washed twice with HBSS, and fluorescence was measured using a plate reader (Tecan) set to excitation wavelengths of 440 and 490 nm. Emission was collected at 535 nm, and the ratio of fluorescence intensities at the two excitation wavelengths was calculated to determine the intracellular pH against a calibration curve which was obtained in parallel with HBSS at pH 6, 7, and 8 supplemented with 5μM of nigericin (Invitrogen).

### Peptide inhibitors of CaMKII

CaMKII peptide inhibitors, custom-made by GenScript (Piscataway, NJ, USA), were designed with sequences identical to those previously used and validated in earlier studies (Ishida et al., 1998; Vest et al., 2007). These include the following myristoyled and unmodified small peptides: CN21, (KRPPKLGQIGRSKRVVIEDDR); scrambled (scr)-CN21, (VKEPRIDGKPVRLRGQKSDRI); AIP-2, (KKKLRRQEAFDAL); and scr-AIP-2, (ELRKFQADLKRKA). Lyophilized trifluoroacetate salts of peptides were resuspended in sterile deionized water to a stock concentration of 1 mM and stored at −20 °C. Some initial experiments were performed with myr-AIP-2 purchased from Calbiochem (ref #189482).

Myristoylated peptides were bath applied at a concentration of 2.5 μM, while unmodified peptides were included in the patch pipette at the same concentration and allowed to diffuse intracellularly for 10 min before running the experiment.

### Proximity ligation assay (PLA)

PLA was performed as previously described (Söderberg et al., 2006; Dinamarca et al., 2016) using cultured cortical neurons plated on glass coverslips, treated at DIV14-17, and fixed for 5 min at 4°C with 4% PFA, 4% sucrose in PBS. Briefly, coverslips were washed 3 times with PBS, permeabilized with 0.1 % Triton X-100 in PBS for 15 minutes, and blocked with 5% BSA in PBS for 30 minutes, all at room temperature. Cells were incubated overnight at 4°C with mouse anti-PSD-95 (1:1000, Abcam ab2723) and rabbit anti-GluN2B (1:1000, Cell Signaling Technology 14544) antibodies in 5% BSA in PBS. After two washes in Wash Buffer A (100 mM Tris-HCl pH 7.4, 150 mM NaCl, 0.05 % Tween20), cells were incubated for 1h at 37°C with secondary antibodies conjugated with oligonucleotides (Duolink probe PLUS and MINUS, Sigma). Coverslips were washed twice in Wash buffer A, and incubated in a ligation solution supplemented with ligase (25 mU/µL) for 30 min at 37°C. Following two rinses in Wash Buffer A, the amplification solution (containing nucleotides and fluorescently labeled oligonucleotides; Sigma) supplemented with polymerase (0.125 U/µL, Sigma) was applied to each sample and incubated for 100 min at 37 °C. To reveal the dendrites of the neurons, coverslips were incubated overnight at 4°C with a mouse anti-MAP2 antibody (1:500, Sigma M4403). After two rinses in Wash buffer A, neurons were incubated for 1 hr at room temperature with Alexa Fluor 488 goat anti-rabbit (1:500, Invitrogen). Finally, coverslips were washed 2 times for 10 minutes in Wash buffer B (200 mM Tris-HC, pH 7.4, 100 mM NaCl), rinsed in 0.05 x Wash buffer B, and mounted on slides with a DAPI containing mounting medium.

### Redox-dependent prediction of intrinsic disorder in GluN2B

Intrinsic disorder scores were calculated according to Erdős and Dosztányi (2020) using the idpr package (version 1.14.0) in R (version 4.4.0) with the mouse GluN2B Uniprot sequence (accession Q01097). The redox-sensitive prediction is done by replacing all cysteine residues to serine when simulating a reducing or “redox-plus” environment (Mészáros et al., 2018).

### Immunoprecipitation

The interaction between CaMKIIα and GluN2B was assessed in synaptosomes from primary cultured cortical neurons by immunoprecipitation. Synaptosomes were isolated using the Syn- PER™ Synaptic Protein Extraction Reagent (ThermoFisher Scientific, 87793) according the manufacturer’s instructions. Protein content was quantified by BCA protein assays and adjusted to a concentration of 1 mg/ml in Pierce IP Lysis/Wash buffer containing 1x Halt™ Protease and Phosphatase Inhibitor Cocktail (ThermoFisher, 78444). Co-IP experiments were performed using Pierce^TM^ Co-Immunoprecipitation kit (ThermoFisher Scientific, 26149) modified to use Dynabeads Protein G (Thermo Fisher, 10004D). Briefly, 150 µg of synaptosome extracts were mixed with 2 µg of either rabbit anti-GluN2B (Abcam, ab65783) or rabbit IgG isotype control antibody (Thermo Fisher 10500C) and incubated under gentle agitation overnight at 4°C to allow the immunocomplex to form. On the next day, 50 µl of Dynabeads were added to the mixture and incubated for 3 h at 4°C. Beads were washed 3 times with 200 µl of Pierce IP Lysis/Wash Buffer. Proteins were eluted from the beads with a 10 min incubation at 95°C in 50 µl of a solution consisting of 2X NuPAGE LDS Sample Buffer (Thermo Fisher, NP0007) and 10% 2-mercaptoethanol. Eluted proteins were diluted 1:1 with water and analyzed using 10% SDS-PAGE Gels.

Antibodies for immunodetection included GluN2B (Invitrogen MA1-2014, 1:1000 diluted in 5% Milk in TBS-T), CaMKIIα (Invitrogen MA1-048, 1:2000 diluted in 5% Milk in TBS-T), PSD-95 (CST 2507, 1:3000 diluted in 3% BSA in TBS-T) or Phospho-GluN2B (EMD Millipore 07-398, 1;3000 diluted in 3% BSA in TBS-T).

### Expression of ANLS/NMDA receptor/CaMKIIα pathway genes in cultured neurons and HEK cells

To quantify the relative expression of genes in cultured cortical neurons and HEK cells, we use RNA-Seq data from one of our previous studies (Margineanu et al., 2018) and from publicly available datasets, respectively. Paired-end reads from untreated neuronal cultures (study SRP150704, NCBI SRA database) and untreated wild-type HEK 293T samples (SRA run accession numbers: SRR3176273, SRR3176274, SRR8181090, SRR8181091, SRR8181092, SRR17873727, SRR17873725, SRR17873726, SRR17873728, SRR22035286, SRR22035287, SRR22035288, SRR17763854, SRR17763855, SRR17763856) were downloaded with fasterq-dump (SRA Toolkit 2.11.3). Adapters and low- quality bases were trimmed with cutadapt, version 18 (Martin, 2011) with the following parameters –q 20 –m 36 – AGATCGGAAGAGCACACGTCTGAACTCCAGTCA -A AGATCGGAAGAGCGTCGTGTAGGGAAAGAGTGT. Reads were then quantified by transcripts indexed on the mouse (neurons) and human (HEK cells) reference transcriptomes (Ensembl release 107 and 108, respectively) using Salmon, version 1.9.0 (Patro et al., 2017), and collated at the gene level in R (version 4.2.1) using the Bioconductor tximeta package (version 1.14.1). Expression levels of genes for the respective cells, normalized for sequencing depth between samples and for gene coding sequence length between transcripts, were expressed as transcript per kilobase million (TPM).

## Results

### Lactate increases the amplitude and the inactivation time constant of NMDAR currents

In previous studies, we demonstrated that lactate upregulates the expression of genes associated with activity and plasticity in an NMDAR-dependent manner (Yang et al., 2014; Margineanu et al., 2018). Additionally, we observed an enhancement of NMDAR signaling linked to NADH formation and the GluN2B subunit’s activity (Yang et al., 2014; Jourdain et al., 2018). Here, intending to gain insights into the molecular mechanism whereby lactate modulates NMDARs, we performed whole-cell patch-clamp recordings of cultured cortical neurons exposed to brief repetitive applications (0.05 Hz) of glutamate (50 μM) and glycine (100 μM) (Figure 1A) in the absence or the presence of 10 mM lactate (Yang et al., 2014; Jourdain et al., 2018). Previous studies have shown that this concentration of lactate exerts robust NMDAR-dependent effects on gene expression and Ca^2+^ signaling (Yang et al., 2014; Jourdain et al., 2018). To isolate NMDAR currents, responses were recorded at different holding voltages in the presence of TTX (0.5 μM) and CNQX (10 μM), blockers of voltage- gated sodium channels and AMPA receptors, respectively. Isolation of NMDAR currents was confirmed by the application D-AP5 (50 μM), a selective NMDAR antagonist, which totally blocked the co-agonist-induced currents (Figure 1B), and by the profile of the I-V relations (Figure 1B), showing characteristics akin to inward rectification due the Mg^2+^ ions block of the channel pore and a reversal of potential around 0 mV (Figure 1C). Lactate presence led to an increased I-V curve slope (Figure 1C) and affected channel inactivation kinetics (Figure 1D).

**Figure 1.**
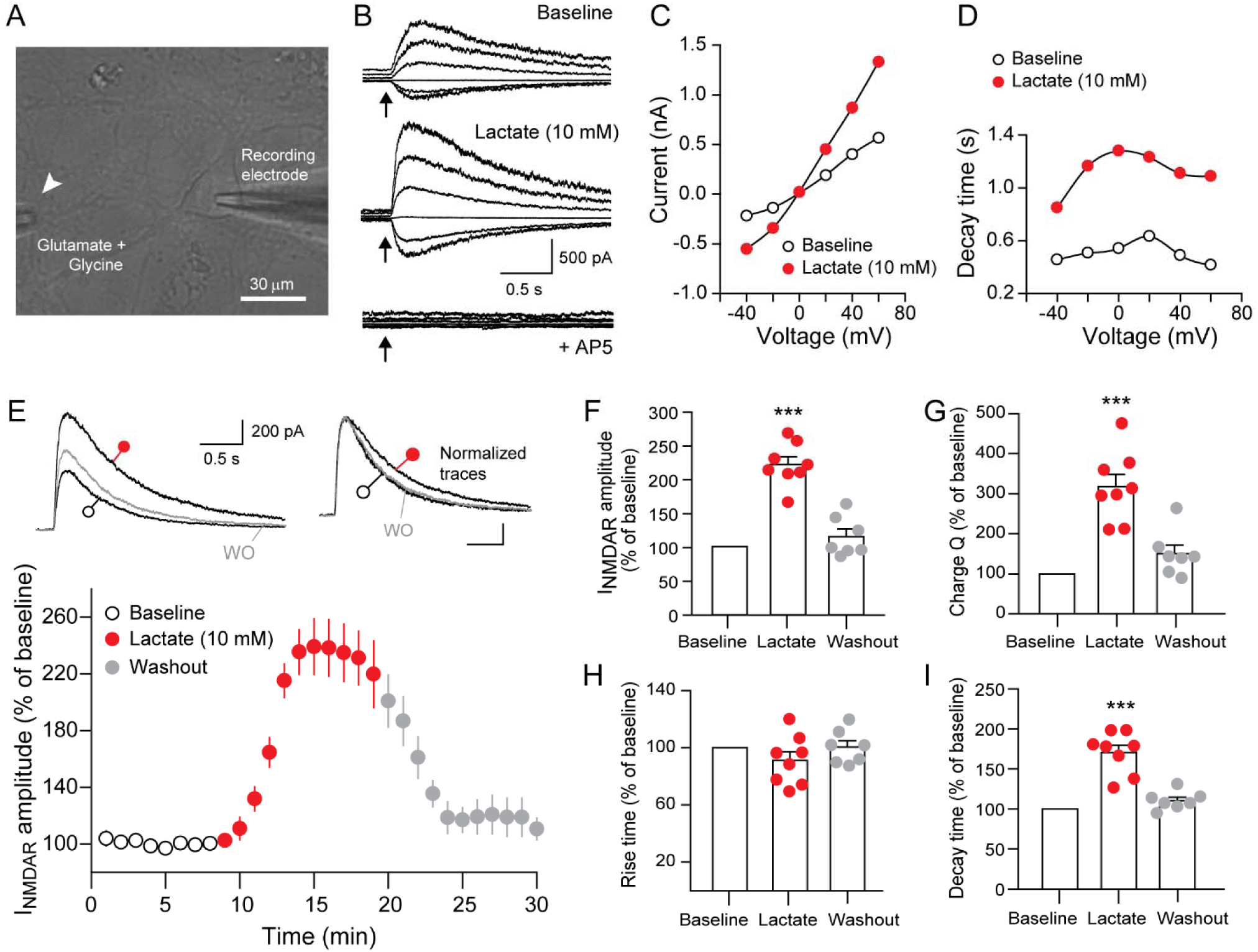
Lactate modulates both the amplitude and the inactivation time constant *τ* of the NMDA receptor current (I_NMDAR_) in cultured cortical neurons. (A) Example of a DIV 14 cultured cortical neuron under patch-clamp, stimulated by brief applications of glutamate (50 µM) and glycine (100 µM) from a micropipette positioned approximately 50 µm from the soma (indicated by a white arrowhead). Scale bar represents 30 µm. (B) Representative isolated I_NMDAR_ traces evoked by puffs of glutamate/glycine at holding voltages ranging from +60 mV to -40 mM (in 20 mV decrement steps) taken at 30 sec intervals during either baseline (top), after 5-6 min in the presence of 10 mM lactate (middle), or in the presence of 10 µM AP5 (bottom). (C) I-V curves and (D) tau values from traces shown in (B). Tau measurements were approximated after fitting the inactivation phase of I_NMDAR_ with a single exponential decay equation using Clampfit software (see Materials and Methods). (E) Top: representative evoked I_NMDAR_ traces recorded at +30 mV before (○), after 5 min of 10 mM lactate treatment (●), or washout (WO, grey traces). Scaled traces (top right) show a slower NMDAR inactivation current in the presence of lactate. Bottom: time-course of averaged normalized I_NMDAR_ peaks before and after 10 mM lactate treatment. Bar charts of normalized values ± SEM of (F) I_NMDAR_ peak amplitude, (G) charge Q, (H) rise time, and (I) decay time during lactate treatment (●) and after washout (●). Measurements were taken 1 min before or 5-6 min after lactate treatment. *** p < 0.001;paired t-test Baseline vs. Lactate.

The lactate-induced potentiation of NMDAR currents, recorded at a holding potential of +30 mV, was rapid, reached a steady-state within 5 minutes, and returned to basal levels upon lactate wash out (Figure 1E). We found that lactate treatment led to robust increases in the peak (Figures 1E and F) and the decay time constant tau (*τ*) (Figures 1E and I) of the NMDARs currents, resulting in a strong augmentation of the charge (Q) transferred (Figure 1G), without affecting the rise time (Figure. 1H). Enhancement of the NMDAR inactivation time constant *τ* was apparent when responses were scaled (Figure 1E, left inset on the top). These findings emphasize lactate’s pivotal role in modulating NMDAR conductance, with potential implications for synaptic function and plasticity.

### GluN2B-containing NMDARs mediates the effects of lactate on the peak responses

NMDARs are tetrameric assemblies composed of two GluN1 subunits paired with two GluN2 (ranging from GluN2A to GluN2D) or GluN3 (GluN3A or GluN3B) subunits (Paoletti et al., 2013). Within the cortical neuronal landscape, GluN1 is the dominant isoform followed by GluN2A and GluN2B (Monyer et al., 1994). The latter is more expressed during development but gives way to GluN2A as neurons progress to maturity (Monyer et al., 1994). This expression profile was corroborated by RNA-seq analysis of our DIV 11 cultured neurons (Figure S1A). NMDARs containing different subunits exhibit distinct physiological properties such as kinetics, leading to variable current flow and different calcium influx upon activation, which may influence neurotransmission and overall plasticity of neuronal networks (Sun et al., 2018; Hansen et al., 2021). To discern a dependence on a specific GluN2 subunit for the potentiation of NMDAR responses by lactate we used PEAQX and ifenprodil, specific inhibitors of GluN2A and GluN2B-containing NMDARs, respectively. In the presence of ifenprodil, lactate was no longer able to potentiate I_NMDAR_ whereas PEAQX did not affect the potentiation by lactate (Figure S1B). These observations indicate that the potentiation of NMDAR currents by lactate was dependent on the GluN2B subunits.

Lactate, traditionally viewed as a metabolic byproduct, is now recognized as a signaling molecule in many organs, including the brain (Magistretti and Allaman, 2018; Certo et al., 2022; Vaccari-Cardoso et al., 2023). Among several pathways, it communicates through a specific G-protein-coupled receptor, HCAR1, which is expressed in various tissues such as adipose tissues and the brain (Liu et al., 2009; Lauritzen et al., 2014; Colucci et al., 2023). Yet, it is unclear if HCAR1 influences lactate’s effects on CNS neurons since its transcripts are barely detectable in cortical neurons (Figure S1A) (Margineanu et al., 2018), and so far reliable tools to study the protein have not been developed (Wallenius et al., 2017; de Castro Abrantes et al., 2019; Briquet et al., 2022). Nevertheless, if HCAR1 is involved in lactate- induced NMDA potentiation, then the specific HCAR1 agonists 3,5-DHBA and 3Cl-HBA should recapitulate the actions of lactate on NMDA currents. Bath application of these drugs at concentrations known to modulate intracellular calcium transients in cortical neurons (Bozzo et al., 2013) neither potentiated the peak I_NMDAR_ amplitude nor altered the inactivation time constant (Figure 2). These observations suggest that lactate acts by entering neurons. In support of this mode of action, we found that pre-incubation of AR-C155858, a selective blocker of monocarboxylate transporters, completely abrogated the lactate-induced increase in the I_NMDAR_ amplitude (Figure 2A and 2B). Interestingly, blocking the cellular uptake of lactate had no impact on the rate of I_NMDAR_ inactivation (Figure 2A and 2C) indicating that lactate affects the amplitude and the decay time by two different mechanisms. Following intracellular accumulation, lactate is transported indirectly via pyruvate or directly into the mitochondria to fuel aerobic respiration (Schurr, 2014; Brooks et al., 2021). First, lactate with NAD^+^ is converted to pyruvate and NADH through a reversible reaction involving predominantly LDH1. Pre-treatment of neurons with the LDH blockers oxamate (6 mM) and stiripentol (200 μM) effectively prevented the lactate-induced potentiation of I_NMDAR_ amplitude (Figure 2A and 2B), yet, it had no discernible impact on the decay time of these currents (Figure 2A and 2C). Compared to lactate, pyruvate elicited a modest increase in the I_NMDAR_ amplitude and a notable effect on the decay time (Figure 2) that paralleled that of lactate, suggesting that a small part of the lactate-mediated effects on NMDAR responses is due to enhanced energy metabolism. In another set of experiments, we exposed neurons to β- hydroxybutyrate (βHB), a ketone body that can substitute neuronal energy needs in the absence of glucose, during periods of fasting, prolonged exercise, or carbohydrate restriction (Jang et al., 2024). βHB was found to also boost both the peak current of NMDAR and prolongs the decay time of its inactivation time constant *τ* (Figure 2). These observations support the notion that the potentiation of I_NMDAR_ amplitudes is specific to monocarboxylates that are oxidized by dehydrogenase using electron acceptors in catabolic pathways while the prolongation of the inactivation time constant *τ* is independent of this pathway and may be common to all monocarboxylates. These collective findings indicate that lactate affects the amplitude and the current inactivation rate by two different mechanisms. The effect on the peak current appears to involve an alteration of the cellular redox balance resulting from the conversion of NAD+ to NADH by the LDH alongside a mild enhancement of cellular respiration, while the effect on the decay time is independent of cellular uptake and lactate oxidation.

**Figure 2.**
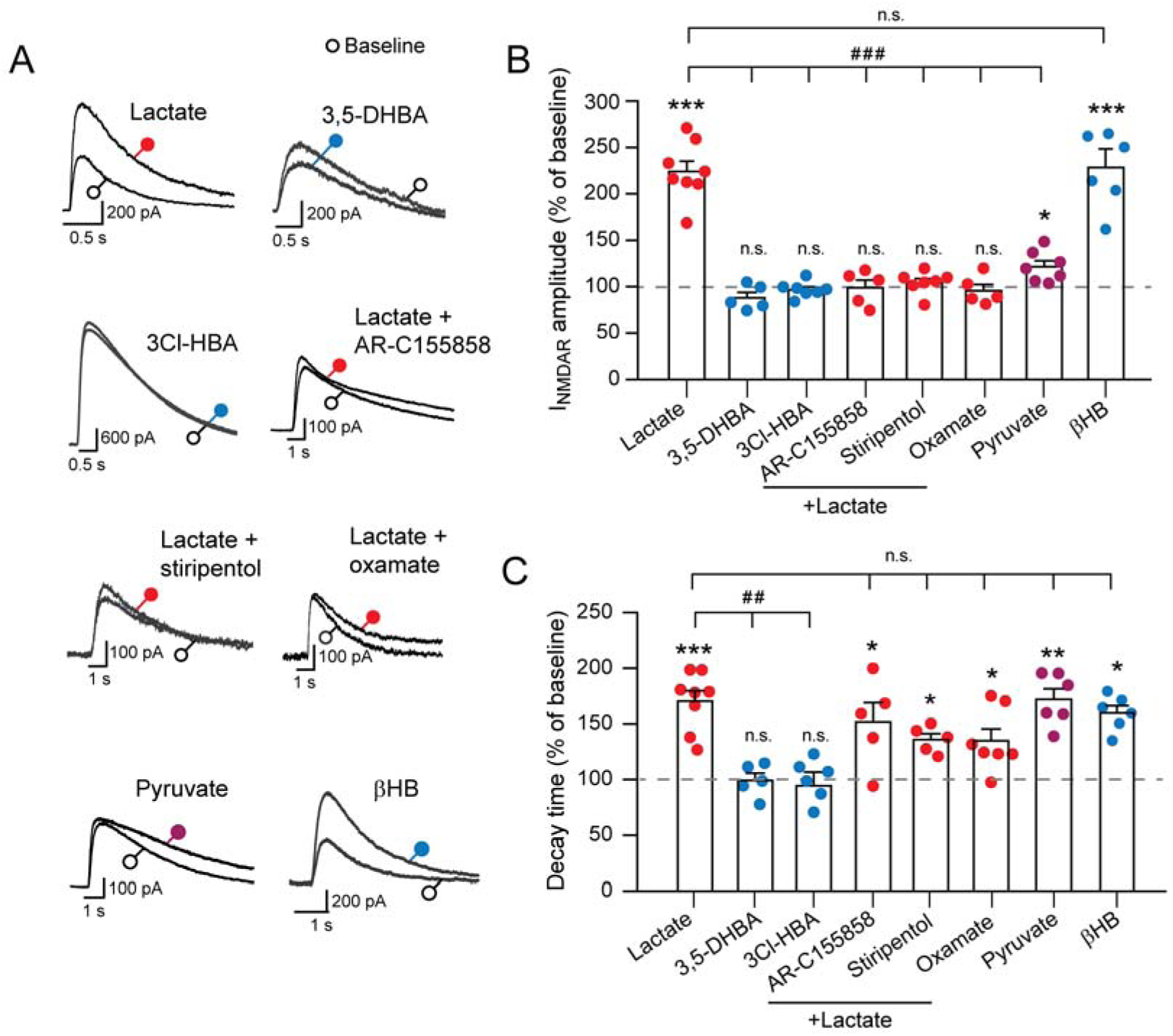
The potentiation of NMDAR currents is specific to lactate and requires cellular uptake and LDH activity. (A) Representative I_NMDAR_ traces evoked by brief applications of NMDAR co-agonists, recorded at a holding potential of +30 mV during the baseline control period (white circles) or treatments (colored circles) with lactate (10mM), HCAR1 agonist 3,5-DHBA (2 mM) or 3Cl- HBA (40 µM), pyruvate (10 mM) or β-hydroxybutyrate (10 mM), with or without AR-C155858 (2 µM), stiripentol (200 µM), or oxamate (6 mM). (B-C) Bar charts summarizing the peak amplitudes (B) and decay times (C) ± SEM of co-agonist-induced I_NMDAR_ in neurons exposed to the conditions described in (A). Statistical significance was determined using paired t-tests between baseline and treatment corrected for multiple comparisons with Tukey’s method (* p < 0.05, ** p < 0.01, *** p < 0.001), and One-Way ANOVA with Holm-Sidak post hoc tests for differences in changes across treatments (# p < 0.05, ## p < 0.01, ### p < 0.001, n.s. = non- significant).

### Critical influence of Ca^2+^ signaling on lactate-induced modulation of I_NMDAR_

There is considerable evidence to suggest that changes in the intracellular redox state can influence Ca^2+^ dynamics and signaling (Zima and Blatter, 2006; Erickson et al., 2008; Requardt et al., 2012; Nikolaienko et al., 2018; Fujii et al., 2023). Moreover, some studies have suggested that lactate has the ability to modulate intracellular levels of calcium. For example, exposure to lactate resulted in calcium elevation in pericytes (Yamanishi et al., 2006), and calcium accumulation is associated with the NMDAR-dependent potentiation of glutamatergic transmission by lactate (Yang et al., 2014; Herrera-López et al., 2020).

However, the mechanism leading to calcium modulation by lactate and whether changes in calcium are a prerequisite or a consequence of lactate’s effects on neuronal activity are unclear. This prompted us to investigate whether changes in intracellular Ca^2+^ dynamics could mediate the effects of lactate on the potentiation of NMDAR function. To this end, whole-cell patch-clamp recordings of co-agonist-induced NMDAR responses acquired at +30 mV from neurons treated with lactate and drugs that interfere with Ca^2+^ signaling were compared to currents obtained in the presence of lactate alone. We found that BAPTA, a potent intracellular calcium chelator, and dantrolene, an inhibitor of ryanodine receptors that mediate calcium release from the endoplasmic reticulum, significantly attenuated the lactate-induced potentiation of peak I_NMDAR_ amplitudes (Figure 3A and 3B). In contrast, treatments with Nisoldipine, an L-type calcium channel blocker, and 2-APB, an IP3 receptor modulator, did not affect the I_NMDAR_ augmentation induced by lactate (Figure 3A and 3B). Further analysis showed that the augmentation of the time constant of the NMDARs by lactate remained largely unaltered across all drug treatments (Figure 3A and 3C. Together, these data suggest that changes in intracellular Ca^2+^ dynamics, resulting in part from ryanodine receptor activation, may shape aspects of the modulation of NMDARs by lactate.

**Figure 3.**
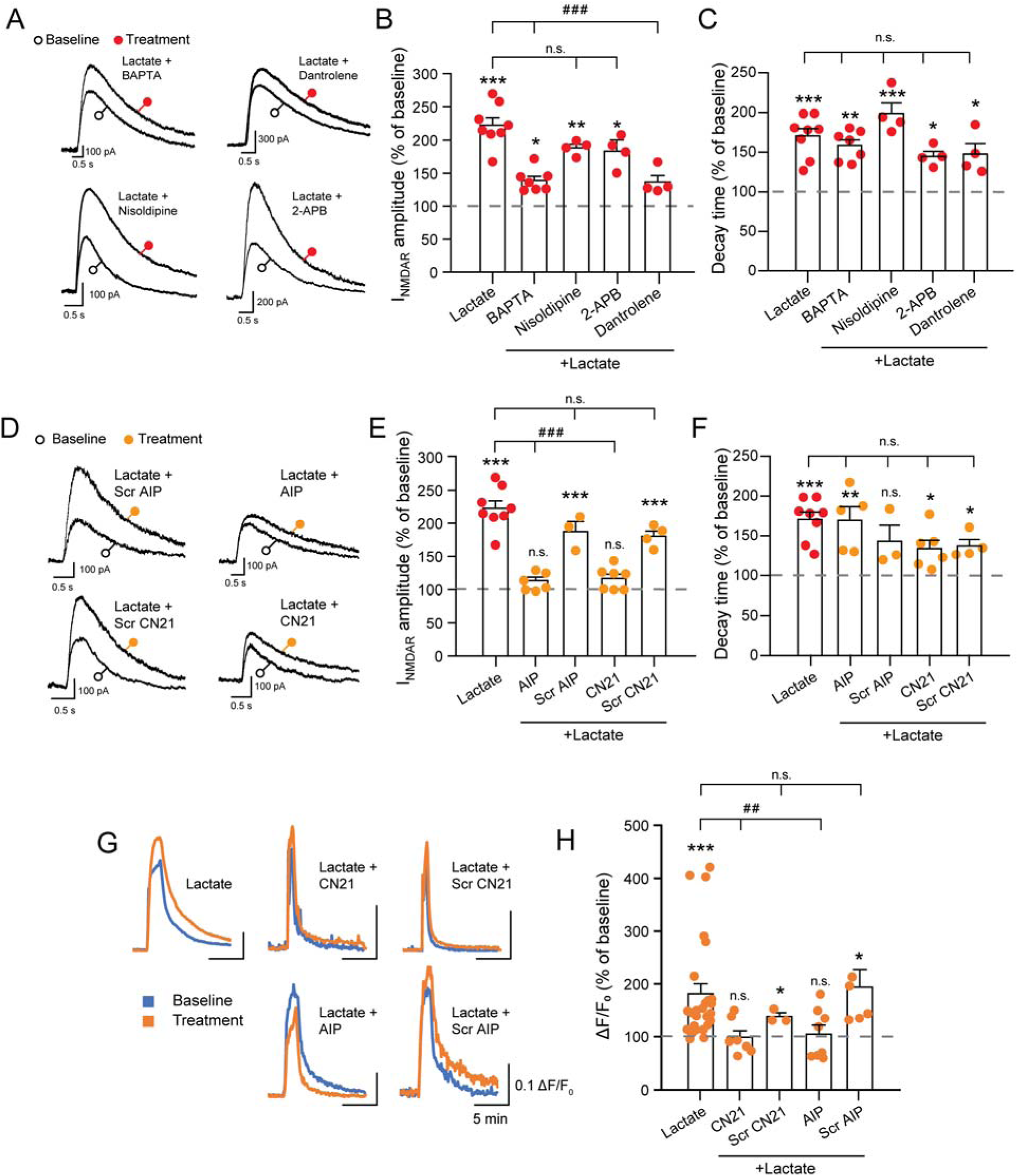
Modulation of NMDAR responses by lactate involves mechanisms and dependency requiring calcium signaling and CaMKII. (A) Representative I_NMDAR_ traces evoked by brief applications of NMDAR co-agonists, recorded at a holding potential of +30 mV during the baseline control period (white circles) or treatments (red circles) with the calcium chelator BAPTA (10 mM), the L-type voltage-gated calcium blocker nisoldipine (2 µM), the IP3 receptor inhibitor 2-APB (50 μM), or the ryanodine receptor antagonist dantrolene (20 µM) in the presence of lactate (10 mM). (B-C) Summary bar charts of I_NMDAR_ amplitudes (B) and decay times (C) recorded in neurons exposed to the calcium modulators as described in (A). (D) Representative I_NMDAR_ traces evoked by brief applications of NMDAR co-agonists, recorded at a holding potential of +30 mV during the baseline control period (white circles) or treatments (oranges circles) with the CaMKII peptide blockers autocamtide-2-related inhibitory peptide (AIP, 10 µM), or CN21 (2.5 µM), along with their respective scrambled (Scr) controls in the presence of lactate. The small peptides were directly introduced to the cytosol via the patch micropipette. (E-F) Summary bar charts of I_NMDAR_ peaks (E) and decay times (F) ± SEM recorded in neurons treated with the CaMKII peptide blockers as described in (D). (G-H) The potentiation of the Ca^2+^ signal by lactate during bath application of NMDAR co-agonists is blocked by CaMKII inhibitors. (G) Superimposed representative intracellular Ca2+ signals expressed as normalized fluorescence changes (ΔF/F_0_) from neurons loaded with Fura-2 following two successive batch applications of glutamate/glycine in the absence (blue traces) or in the presence of 10 mM lactate (orange traces) in cultures exposed to the myristoylated-CaMKII peptide blockers AIP (10 µM), CN21 (2.5 µM), or their respective scrambled (Scr) controls (10 µM). (H) Bar chart summary of lactate’s potentiating effects on Ca^2+^ signals. Data are expressed as percentages ± SEM of the normalized fluorescence changes (ΔF/F_0_) in the presence of lactate over responses without lactate in neurons exposed to CaMKII peptide blockers and their respective scrambled controls. Statistical significance was determined as described in Figure 2.

Given the central role of calcium/calmodulin activated CaMKII in synaptic plasticity and its known interactions with NMDARs (Bayer and Schulman, 2019; Yasuda et al., 2022; Nicoll and Schulman, 2023), it is plausible that CaMKII may also be a key player in the lactate-induced potentiation of these receptors. Utilizing patch-clamp recordings and calcium imaging, we examined NMDAR responses induced by puffs of glutamate and glycine, with or without the application of the CaMKII small peptide inhibitors AIP2 and CN21, which both block the kinase activity and disrupt GluN2B binding (Pellicena and Schulman, 2014; Yasuda et al., 2022; Nicoll and Schulman, 2023). Our results, as illustrated in Figure 3D and 3E, demonstrated that the infusion of AIP2 or CN21 via the patch pipette markedly abrogated the potentiation by lactate of peak NMDAR responses. In contrast, blocking CaMKII had modest effects on the increase of the inactivation time constant of NMDARs by lactate (Figure 3D and 3F). Similarly to the electrophysiological readout, the augmentation of the calcium indicator fura-2 signal induced by 2 min application of 1 µM of glutamate and 100 µM of glycine (Jourdain et al., 2018) in the presence of lactate was eliminated by bath application of membrane permeant versions of these peptides (Figure 3G and 3H). Absence of effect of the scrambled peptides in these calcium imaging experiments confirmed the specificity of the CaMKII inhibitory peptides in blocking the lactate effects (Figure 3G and 3H). These findings using complementary approaches demonstrate that the recruitment of CaMKII is necessary for the potentiation of NMDAR by lactate.

### Lactate Potentiates NMDARs via CaMKII and GluN2B Interaction in HEK Cells

In order to better understand the role of CaMKII on the potentiation of NMDAR by lactate, we took advantage of a well-established pharmaco-genetic cellular model afforded by HEK cells, which do not express CaMKIIα (Figure 2SD and 2SE) nor NMDAR subunits (Figure S2A and S2D). These versatile and readily transfectable cells are well suited to study the NMDAR responses originating from the combination of NMDAR subunit variants and meet the criteria to assess the actions of lactate (Figure S2B and S2C). To evaluate the potentiation of NMDAR responses, we performed electrophysiological recordings of HEK cells stably expressing CaMKIIα (CaMKIIα) or not (WT) transiently transfected with GluN1 and GluN2B expressing constructs. First, we tested whether lactate alters the conductance of NMDAR responses. In these experiments, current-voltage relationships were generated by stepping the membrane potential of GluN1 and GluN2B transfected cells from -30 mV to +50 mV while delivering brief pulses of glutamate and glycine with a micropipette located at 40-60 μM from the cell body in the absence or presence of lactate. Figures 4A and 4B show typical J-shaped current-voltage relationships for NMDARs in the presence of magnesium ions that are blocked by AP5. In WT HEK cells, lactate exposure resulted in no discernible change in the I- V curve profiles (Figure 4B, left panels). In contrast, in CaMKIIα HEK cells, lactate steepened the I-V curves, indicating enhanced conductance compared to the control condition (Figure 4B, right panels). Notably, lactate increased to the same extent the decay time of NMDAR inactivation currents in both the WT and the CaMKIIα-expressing HEK cells (Figure 4B, bottom panels). Quantification at a holding potential of +30 mV revealed that lactate but not pyruvate amplified NMDAR current amplitudes in the presence of CaMKII (Figure 4C top), while the two monocarboxylates independently of CaMKII prolonged the decay time (Figure 4C, bottom). Additionally, the peak I_NMDAR_ potentiation by lactate in HEK cells expressing CaMKIIα was fully inhibited by the MCT inhibitor AR-C 155858 (Figure S3A) but not the decay time, indicating that the cellular uptake of lactate is also required in HEK cells. Collectively, these findings demonstrate that CaMKII is essential for lactate-induced potentiation of NMDARs.

**Figure 4.**
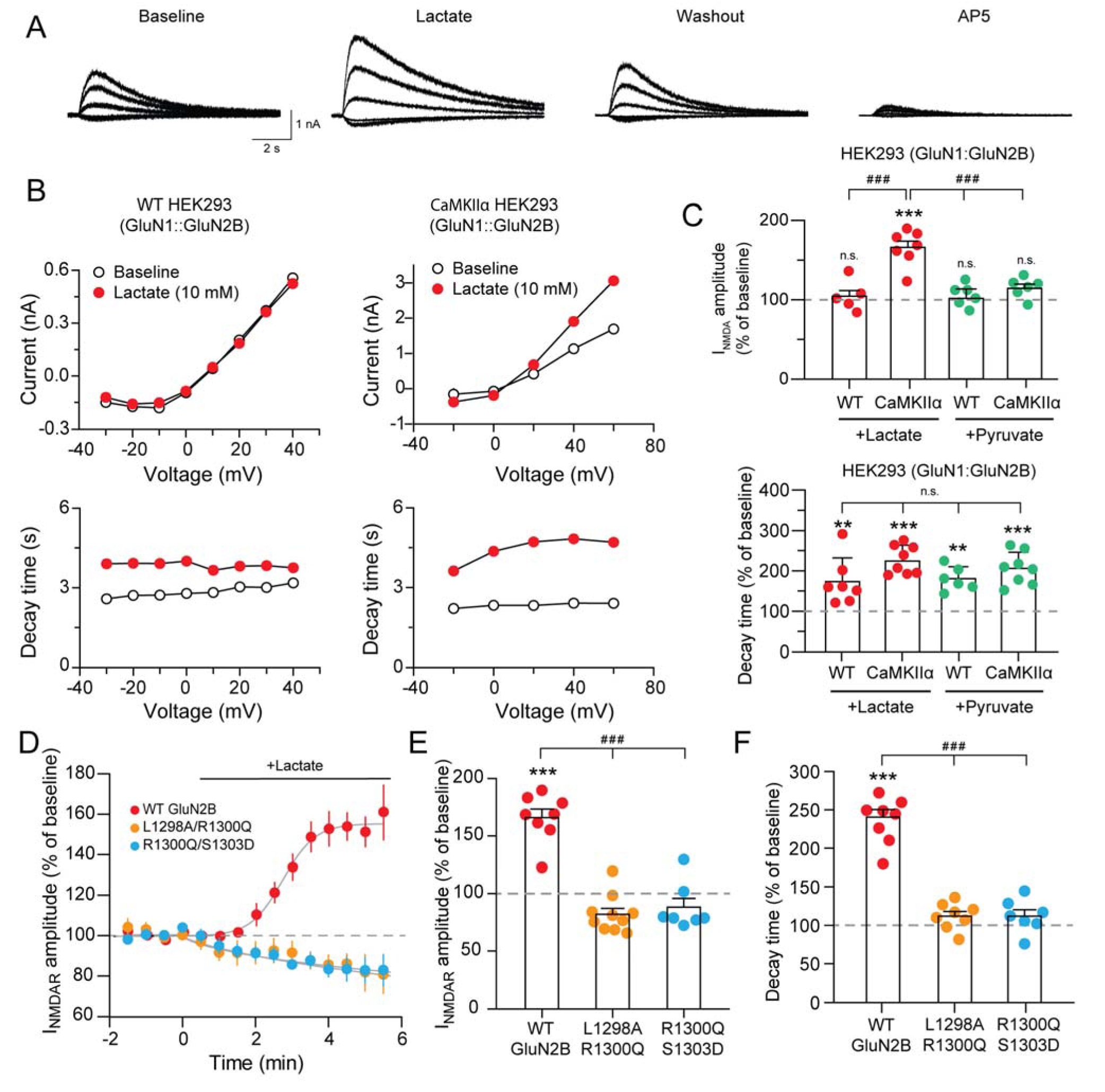
Lactate-induced NMDAR potentiation in HEK cells hinges on the expression of CaMKII and its interaction with the GluN2B subunit. (A) Representative isolated I_NMDAR_ traces recorded from CaMKII-expressing HEK cells transiently transfected with expressing constructs encoding the NMDAR subunits GluN1 and Glu2B. Currents were evoked by puffs of glutamate/glycine at holding voltages ranging from +60 mV to -20 mM (in 20 mV decrement steps), recorded at 30-sec intervals during baseline (control), after 5-6 min in the presence of 10 mM lactate, during the washout period, or in the presence of 50 µM AP5. (B) Lactate stimulates peak I_NMDAR_ in HEK cells expressing functional NMDAR (GluN1::GluN2B) in CaMKIIα-expressing HEK cells (right panels) but not in cells lacking CaMKIIα (WT, left panels). This effect is noticeable at holding potentials above 20 mV. In contrast, lactate prolongs decay times in both types of HEK cells, independent of CaMKIIα expression (bottom panels). (C) Summary bar charts of I_NMDAR_ amplitudes (top) and decay times (bottom) ± SEM in response to puffs of co-agonists at a holding potential of +30 mV, in the presence of lactate or pyruvate in NMDAR-expressing HEK cells with or without CaMKIIα expression. (D) Time-course of peak I_NMDAR_ in the absence and presence of 10 mM lactate for CaMKIIα-expressing HEK cells transfected with GluN1 and either WT or mutant GluN2B subunits. The two GluN2B variants (L1298/R1300Q and R1300Q/S1203D) are known to disrupt the interaction between GluN2B and CaMKIIα. (E, F) Quantitative summaries of the I_NMDAR_ amplitudes (E) and decay times (F). Bar charts are expressed as percentages of averaged baseline values ± SEM. Statistical significance was determined as described in Figure 2.

CaMKII is known to interact with the GluN2 subunits of NMDARs (Gardoni et al., 1998; Strack et al., 2000; Bayer et al., 2001; Zhou et al., 2007; O’Leary et al., 2011; Halt et al., 2012). This interaction is a pivotal component in synaptic plasticity, particularly in long-term potentiation (LTP). We therefore explored the possibility that the binding between CaMKII and GluN2B was necessary for the potentiation of NMDAR function by lactate. Previous mutational studies demonstrated that CaMKII, through its catalytic domain including the S and T sites (Özden et al., 2022), binds to the region around Ser1303 of GluN2B (Leonard et al., 1999; Strack et al., 2000; Bayer et al., 2001; Mayadevi et al., 2002). To examine the functional role of the complex formation, we used two different dual GluN2B mutants (R1300Q/S1303 and L1298A/S1303) known to interfere with the binding between GluN2B and CaMKII (Strack et al., 2000; Halt et al., 2012). Co-transfection of these mutants with GluN1 in CaMKIIα- expressing HEK cells resulted in functional expression of NMDAR showing responses to brief applications of glutamate and glycine similar to those with the wild-type subunits (Figure S3C). However, in CaMKIIα-expressing HEK cells transfected with either of these mutants, lactate was not effective in potentiating the co-agonist-induced I_NMDAR_ (Figure 4 D-F). The time course and summary of the normalized I_NMDAR_ amplitudes recorded at +30 mV before and after the application of lactate shows that the L1298A/R1300Q or the R1300Q/S1303D mutant GluN2B completely abolished the increase seen with the unmutated GluN2B (Figure 4A), while the mutants did not significantly affect the lactate-induced changes in the I_NMDAR_ decay times recorded at 5 min (Figure 4C). These data indicate that the binding between CaMKIIα and GluN2B is required for the potentiation of NMDAR responses by lactate.

### Molecular mechanism for the amplification of NMDAR currents by lactate-induced changes in the intracellular redox state

Exposing neurons to altered redox environments is well known to modulate NMDAR responses (Kohr et al., 1994; Sullivan et al., 1994; Choi and Lipton, 2000), possibly through changes in the redox state of extracellular cysteine residues in the NMDAR (Sullivan et al., 1994; Lipton et al., 2002). It is however less clear whether changes in the intracellular redox balance can modulate NMDAR function. We found that introducing NADH via the patch pipette delayed by several minutes the time course of the potentiation of the I_NMDAR_ amplitudes by lactate (Figure 5A). This observation suggests that perturbation of the intracellular redox ratio upon lactate entry may regulate the magnitude of the NMDAR responses. Figure 5B supports this hypothesis, showing that the intracellular infusion of the reducing agents dithiothreitol (DTT) and β-mercaptoethanol (βME), alongside lactate exposure, does not alter the magnitude of the lactate-induced potentiation. This indicates that lactate and the reducing agents act through the same mechanism. Conversely, the presence of the oxidizing chemical 5,5-dithio-(bis-2-nitrobenzoic) acid (DTNB) in the patch pipette abolished the lactate-induced potentiation, bringing the current amplitude back to baseline levels. These results suggest that the potentiation of NMDAR currents is modulated by an amplification of the reduced intracellular environment induced by lactate.

**Figure 5.**
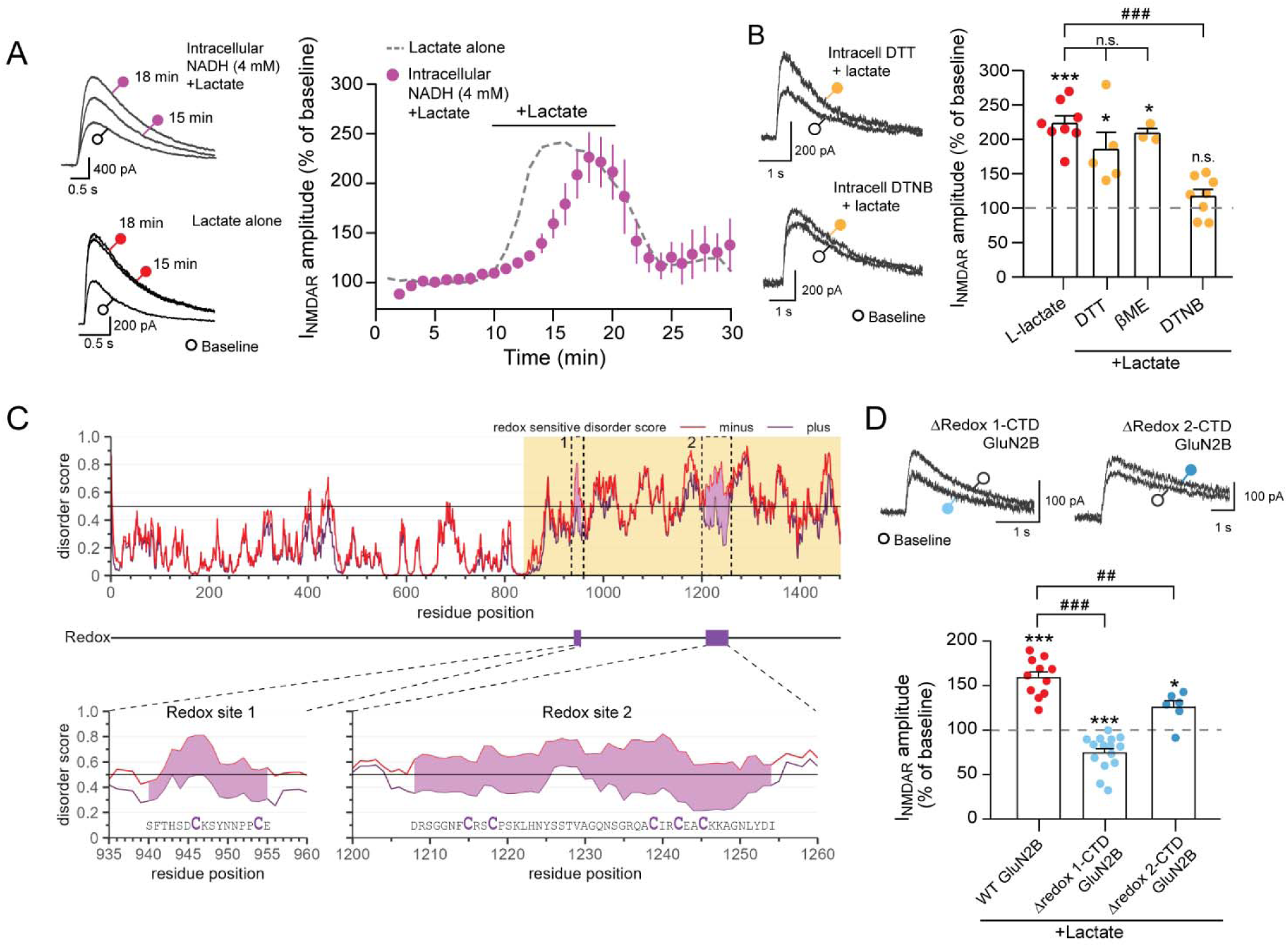
Lactate potentiation of NMDAR responses depends on two redox-sensitive cysteine-rich sequences in the intrinsically disordered CTD of GluN2B subunit (A, left panels) Representative traces of I_NMDAR_ evoked by puffs of glutamate/glycine at the soma, recorded in cultured neurons at a holding potential of +30 mV. Traces during the baseline period are marked with a white circle, and those 5 and 8 minutes after the onset of the lactate treatment with a colored circle. Top example traces include 4 mM NADH in the intracellular patch pipette solution, while bottom traces do not. The representative traces with lactate were recorded 15 and 18 minutes after going whole-cell. (A, right panel) The time- course of I_NMDAR_ amplitudes induced by co-agonists is shown before, during, and after lactate exposure, with (purple dots) and without (dashed line) intracellular NADH. The dashed line illustrates the time-course of I_NMDAR_ amplitudes without NADH, as shown in Figure 1E. (B, left panels) Representative I_NMDAR_ traces evoked by brief applications of NMDAR co-agonists were recorded in cultured neurons at a holding potential of +30 mV with a patch pipette solution containing the reductant DTT (top) or the oxidizer DTNB (bottom) during treatment with lactate (orange circle) or the baseline control period (white circle). (B, right panel) Bar chart summarizing the peak I_NMDAR_ responses ± SEM in neurons exposed to lactate (10mM) with or without intracellular infusion of reductants (DTT, 1 mM and βME, 300 µM) or the DNTB (200 µM). (C) Prediction of intrinsic disorder by residue position in mouse GluN2B in reducing (plus, dark purple line) or oxidizing/normal (minus, red line) conditions. Redox- sensitive stretches are identified by the divergence of the red and dark purple lines across the disorder score threshold of 0.5. These regions are highlighted in light purple and marked by dashed rectangles that form areas highlighted in light purple delineated by dashed rectangles. The beige background indicates the C-terminal domain (CTD) of the protein. (D, top) Representative I_NMDAR_ traces evoked by brief applications of NMDAR co-agonists, recorded at a holding potential of +30 mV in HEK cells in the presence of lactate (colored circles) or during the baseline control period (white circle). (D) Quantitative summaries of the effect of 10 mM lactate (expressed as percentage of baseline) on I_NMDAR_ amplitudes in CaMKIIα- expressing HEK cells transfected with GluN1 and WT Glu2B (WT), C946-C954S mutant GluN2B (Δredox 1-CTD GluN2B), or C1215S-C1218S-C1239-C1242S-C1245S GluN2B (Δredox 2-CTD GluN2B) subunits. Summary data are expressed as percentages of averaged baseline values ± SEM. Statistical significance was determined as described in Figure 2.

It is known that reducing and oxidizing agents act by modifying proteins containing thiol groups (cysteines and methionines). These post-translational modifications have been extensively documented for their role as a signaling mechanism (Paulsen and Carroll, 2013). With this in mind, we hypothesized that lactate could act through this form of signaling by reducing thiol-containing proteins involved in NMDAR signaling.

CaMKII is known to undergo reversible oxidation of cysteines and methionines, which, like autophosphorylation, sustains its activity once Ca^2+^ is no longer bound to CaM (Erickson et al., 2011; Coultrap and Bayer, 2014; Rocco-Machado et al., 2022). To test whether lactate- induced changes in the intracellular redox state affect NMDAR response by modulating the redox state of key residues in CaMKII’s regulatory domain, we examined the effect of lactate on NMDAR potentiation in HEK cells expressing redox-insensitive CaMKII (Figure S4A and see Materials and Methods). As shown in Figure S4B, lactate potentiated NMDAR currents in a similar fashion in both redox-insensitive (Δredox) and wild-type CaMKII-expressing HEK cells. This indicates that lactate does not influence the redox state of these key residues on CaMKII or that modification of their redox state is not crucial for lactate’s effect on NMDAR potentiation.

The C-terminal domain (CTD) of GluN2B is known to include large portions of intrinsic disorder (Choi et al., 2011) and is enriched in cysteine residues. These features may play a role in redox chemistry and protein functionality (Choi et al., 2013). To investigate whether intracellular redox changes, such as those caused by lactate entry and oxidation to pyruvate, could affect NMDAR structure, we predicted the disordered regions of the CTD in different redox states (Figure 5C). We identified two potentially redox-sensitive cysteine-rich sequences within the disordered CTD. The first sequence, spanning residues 940-955, contains two cysteines. The second, longer sequence, between residues 1208 and 1254, includes five cysteines. These regions, indicated in Figure 5C by purple shadings marking the separation in the disorder scores, show sensitivity to redox changes, highlighting their potential role in NMDAR regulation.

To investigate the role of redox-sensitive regions in the CaMKII-dependent regulation of NMDAR by lactate, we generated two redox-insensitive GluN2B mutants (Δredox 1-CTD and Δredox 2-CTD, see Materials and Methods and Figure 5C) based on the modeling results where cysteines were replaced with serines. Co-transfection of these mutants with GluN1 in CaMKIIα-expressing HEK cells resulted in functional NMDARs that responded to brief applications of glutamate and glycine similarly to wild-type subunits in baseline conditions (Figure 5D, top). However, in CaMKIIα-expressing HEK cells transfected with the Δ redox 1- CTD GluN2B mutant, lactate failed to potentiate the co-agonist-induced I_NMDAR_ (Figure 5D). The Δ redox 2-CTD GluN2B mutant also exhibited diminished potentiation by lactate, although to a lesser extent than the Δ redox 1-CTD mutant (Figure 5D). These results suggest that redox modifications of the CTD of NMDAR are crucial for lactate-induced potentiation of NMDAR.

To further elucidate the role of the CaMKII/NMDAR complex in synaptic plasticity changes by lactate in neurons, we first conducted a series of quantitative protein assays in neurons treated with pyruvate or lactate. For these experiments, we employed more mature cultured cortical neurons (DIV 14-17) rich in synaptic contacts. Exposure of these neurons to 10 mM lactate or pyruvate for 20 min did not affect the CaMKII or GluN2B protein expression levels in neurons or in the synaptosome-enriched fractions (Figure 6A, and Figure S5B and C).

**Figure 6.**
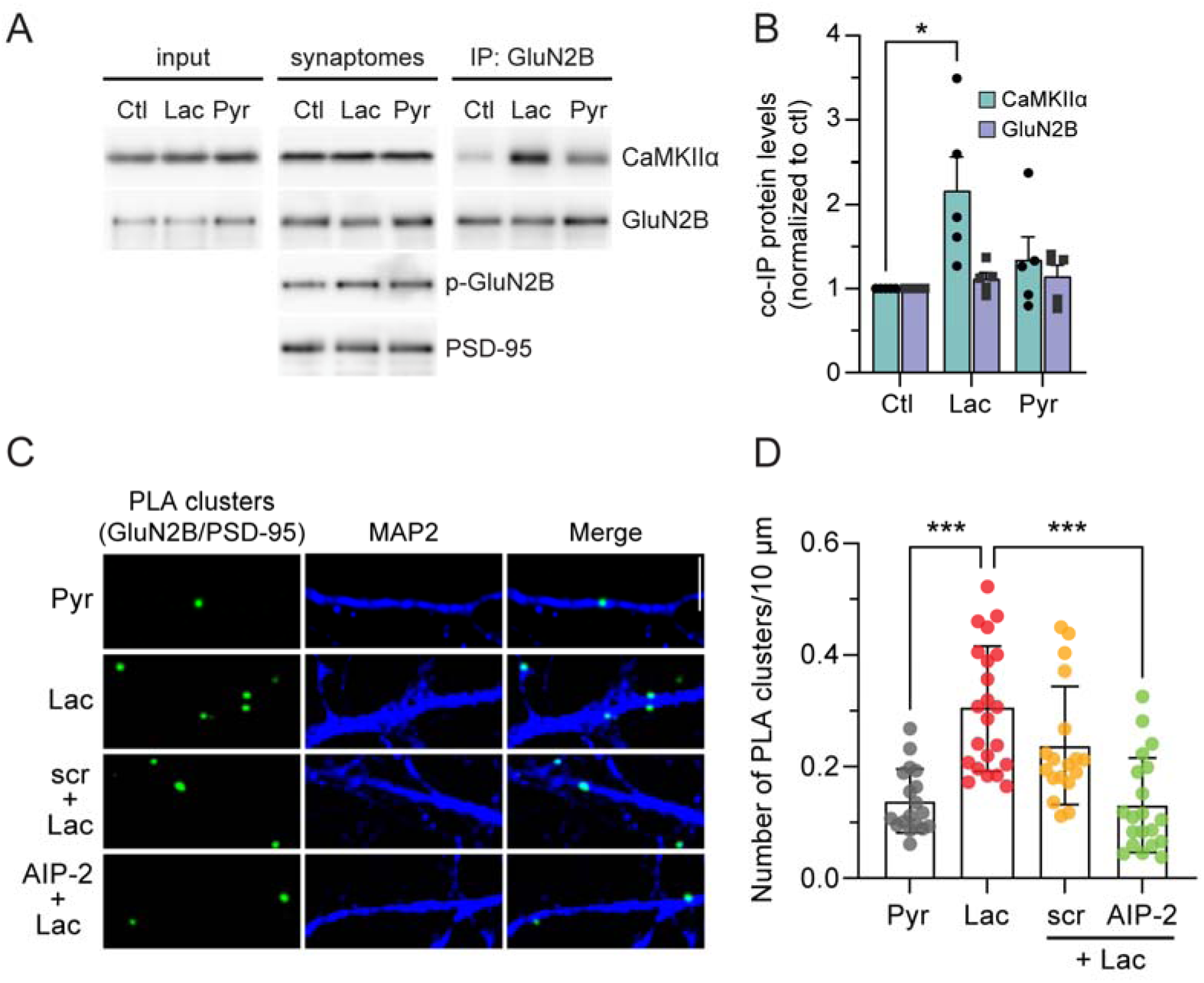
Lactate potentiates the binding between CaMKII and GluN2B and drives a CaMKII-dependent increase of GluN2B-PSD-95 clustering. (A) Lactate increases the association between GluN2B and CaMKII. Representative western blots showing expression of CaMKII and GluN2B in total protein extract (input), CaMKII, GluN2B, p1303-GluN2B, and PSD-95 in synaptosome extract (synaptosomes), and CaMKII and GluN2B in synaptosome extracts immunoprecipitated with GluN2B antibody (IP: GluN2B) treated or not with 10 mM of lactate (Lac) or pyruvate (Pyr). (B) Quantitative summary plot for the synaptosome-enriched proteins immunoprecipitated with a GluN2B antibody expressed as a ratio of their respective control (Ctl) ± SEM. Statistical significance: * p < 0.05, Two-Way ANOVA followed by Dunnett’s multiple comparisons test. (C) Representative images of proximity ligation assays (PLA) for GluN2B/PSD-95 (green) in cultured cortical neurons treated with pyruvate (Pyr, 10 mM) or lactate (Lac, 10 mM) in the presence or absence of the CaMKII inhibitor peptide myr-AIP-2 (1 µM) or its scrambled control (scr, 1 µM). MAP2 immunostaining (blue) was used to delineate the dendrites. Scale bar, 5 µM. (D) Quantitative summary with data expressed as the mean ± SD. PLA cluster numbers per 10 µM of dendrite. ***p<0.001, One-Way ANOVA followed by Bonferroni’s multiple comparison tests.

Moreover, these treatments did not increase the levels of phospho-Ser1303-Glu2B in the synaptosome fractions (Figure 6A and Figure 5C), a post-translational modification induced by CaMKII (Omkumar et al., 1996) known for shaping NMDAR functionality linked to synaptic plasticity (Nicoll and Schulman, 2023) or those of the postsynaptic density marker PSD-95 (Figure 6A and Figure 5C). These data suggest that the monocarboxylates do not affect the activity of CaMKII toward its substrate GluN2B, nor do they alter the expression levels of the main postsynaptic density marker. Co-immunoprecipitation assays were conducted next to investigate the quantitative interaction between CaMKII and GluN2B in synaptosomal fractions derived from these cultures. As an initial step to ensure specificity of the observed interactions, we performed control experiments using control IgG or omitting the IgG entirely (Figure S5A). Upon treating cultured neurons with lactate, we found a marked increase in the co-IP of CaMKII with GluN2B (Figure 6A and 6B). This enhancement in interaction was notably greater than what was observed when neurons were treated with pyruvate (Figure 6A and 6B), suggesting a specific modulatory role for lactate in promoting the association between CaMKII and GluN2B at synapses.

Finally, we performed proximity ligation assays (PLA) to examine whether lactate signaling could influence the postsynaptic enrichment of GluN2B-containing NMDARs. As shown in Figure 6C and D, PLA experiments with antibodies against GluN2B and the postsynaptic marker PSD-95 revealed that, in comparison to pyruvate-exposed cultures, used here as an equi-caloric control, when neurons were treated with lactate for 20 min, there was a significant increase in the number of PLA clusters. This observation suggests that lactate signaling enhances the prevalence of synapses where GluN2B is expressed in close proximity to PSD-95. Moreover, the lactate-induced augmentation in the PLA cluster count was fully prevented by pre-treating the cells with the CaMKII inhibitory peptide myr-AIP, while a scrambled version of this inhibitory peptide had no effect on the observed outcome (Figure 6C and 6D). Together, these data underscore the requirement and the specificity of CaMKII’s role in modulating the proximity for interaction between GluN2B and PSD-95 in response to lactate exposure.

## Discussion

Lactate has emerged over the last decade as not only an energy substrate intrinsically linked to neuronal activity but also as a signaling molecule in particular for synaptic plasticity (Barros, 2013; Magistretti and Allaman, 2018). Through the set of experiments presented in this article we wanted to understand the molecular mechanisms that link the behavioral effects of lactate on synaptic plasticity, learning and memory with the enhancement of highly regulated NMDAR currents that underlie synaptic plasticity.

Our findings demonstrate that lactate enhances NMDAR responses by increasing both the amplitude and inactivation time constant of NMDAR currents in cultured cortical neurons. These effects are specifically driven by NMDARs containing GluN2B subunits, the most prevalent forebrain NMDAR subtype playing a crucial role in neuronal development and synaptic plasticity (Paoletti et al., 2013; Ge and Wang, 2023). The mechanism of this amplification of NMDAR currents involves the uptake of lactate into neurons and its metabolic conversion to pyruvate, concomitantly converting NAD+ to NADH, a reaction that modifies the cellular redox state and, in turn, stimulate NMDAR. We found that this redox homeostasis shift promotes Ca^2+^ mobilization from the endoplasmic reticulum and creates an environment that affects the CTD of NMDAR via cysteine modification. This change favors the interaction between CaMKII and GluN2B subunits, facilitating the recruitment of NMDARs to the synapse and boosting synaptic plasticity. To better illustrate the complex cellular pathways and molecular interactions involved in lactate’s role as both an energy substrate and a signaling molecule, we have summarized our findings and working hypothesis in a schematic representation shown in Figure 7.

**Figure 7.**
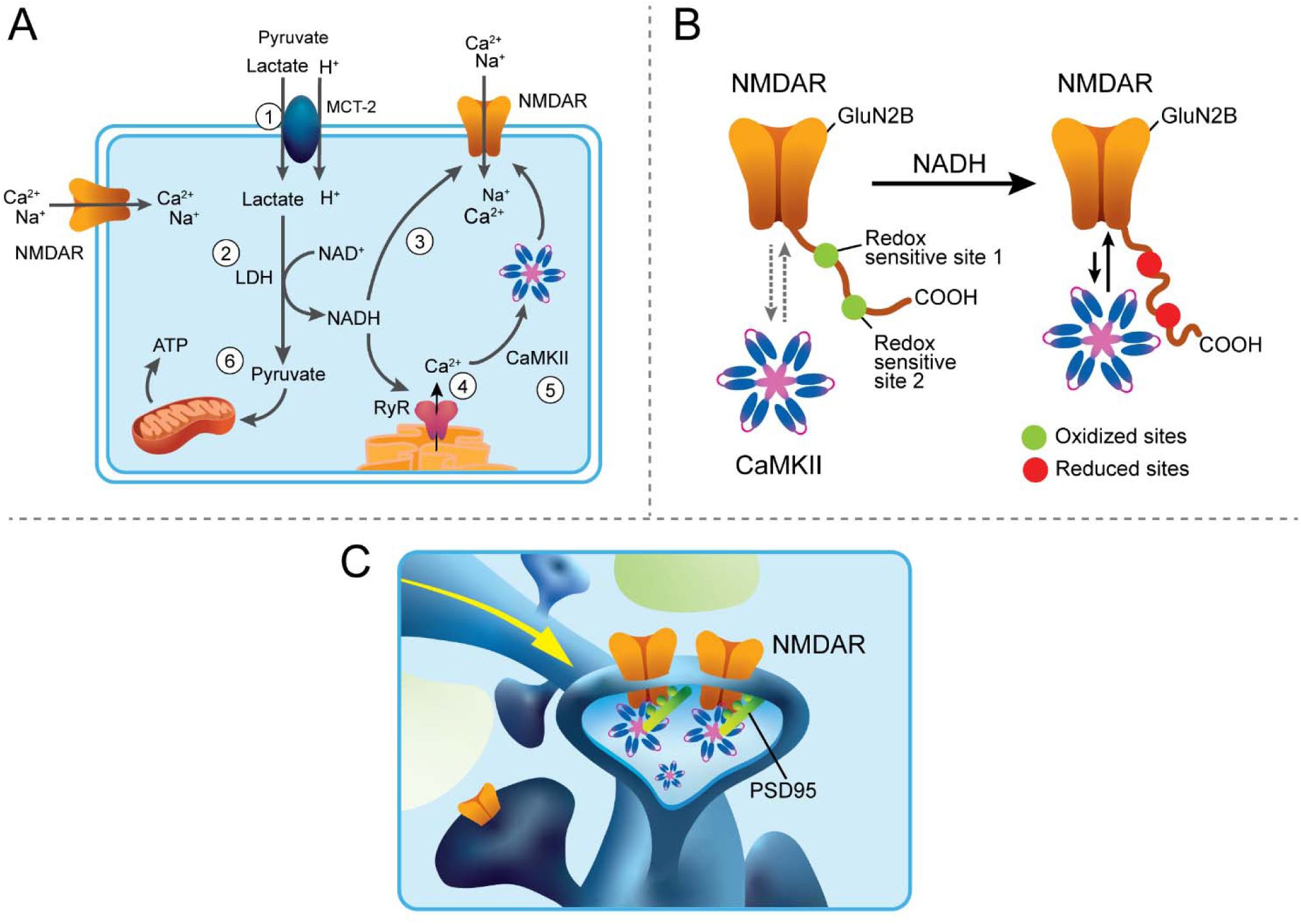
Schematic representation of the cellular mechanisms involved in the modulation of NMDAR responses and facilitation of synaptic plasticity by lactate. (A) Extracellular lactate is transported passively along its concentration gradient through the monocarboxylate transporters 2 (MCT-2) into neurons (1) where it is converted by the lactate dehydrogenase (LDH) in presence of the oxidized form of nicotinamide adenine dinucleotide (NAD^+^) to pyruvate and the reduced form of nicotinamide adenine dinucleotide (NADH) (2). This metabolic process increases the NADH/NAD+ ratio, leading to a more reduced intracellular milieu. Such a change in the internal redox state, in turn, affects cysteine-rich redox-sensitive sites on the C-terminal domain of the NMDAR GluN2B subunits favoring CaMKII binding (3) and modulates the activity of ryanodine receptors (RyRs), causing, together with the activation of NMDAR, an increase in intracellular Ca^2+^ levels (4). Elevated intracellular Ca^2+^ levels activate CaMKII which allow its binding to NMDAR (5). Pyruvate is imported into the mitochondria and metabolized by the tricyclic acid cycle and the oxidative phosphorylation to produce ATP to sustain the energy needs of neurons (6). (B) A detailed view of the molecular interaction between NMDARs and CaMKII under the influence of lactate. Lactate-induced NADH production alters the intracellular redox environment, promoting the rearrangement of the intrinsically disordered structure of redox-sensitive cysteine-rich sites within the C-terminal domain of GluN2B. These modifications, along with the activation of CaMKII by Ca^2+^, promote the binding of CaMKII to NMDARs, stimulating the accumulation of NMDAR at synapses, illustrated by an increase in the association between GluN2B and PSD-95, ultimately enhancing synaptic NMDAR function and synaptic plasticity (C).

Most importantly we found that the establishment of a reduced environment by the conversion of lactate to pyruvate targets two redox-sensitive cysteine-rich stretches in the intracellular C- terminal domain of the GluN2B subunit of NMDARs. Interestingly, the CTD of the GluN2B subunit of the NMDAR is intrinsically disordered (Choi et al., 2011; Warnet et al., 2021) and contains many cysteine residues. In some conditionally disordered proteins, changes of the oxidation status are coupled to disorder-to-order or order-to-disorder transitions (Reichmann and Jakob, 2013) with cysteine residues undergoing reversible thiol oxidation to dynamically respond to the redox status of the environment. Changes in the intrinsic order may facilitate versatile interactions with multiple proteins and play critical roles in signaling pathways (Wright and Dyson, 2015; Kjaergaard and Kragelund, 2017). This characteristic enables intrinsically disordered regions to participate dynamically in numerous cellular processes by providing binding sites for various signaling and scaffold proteins. Thus, the NADH-mediated lactate-induced changes in the intrinsic disorder of the CTD of GluN2B is likely responsible for the enhanced interaction between the GluN2B and CAMKII. Recent studies have shown that the maintenance of long-term potentiation (LTP) primarily relies on the structural, rather than the enzymatic, role of CaMKII, particularly its interaction with GluN2B (Incontro et al., 2018; Tullis et al., 2023; Chen et al., 2024). The remodeling of the CTD of GluN2B through lactate-induced redox changes could thus potentiate this structural interaction facilitating LTP.

Furthermore, the interaction between CaMKII and GluN2B is essential for the formation of phase-separated protein condensates at the synapse (Hosokawa et al., 2021), which serve as organizing hubs for synaptic signaling associated with synaptic plasticity (Wu et al., 2020; Nicoll and Schulman, 2023). By remodeling the CTD of GluN2B, lactate-induced redox changes could enhance the stability or formation of these condensates, thus promoting the structural role of CaMKII at the synapse. This interaction favored by lactate may ensure the persistence of synaptic changes by maintaining the structural integrity of the synaptic architecture and facilitating the recruitment of additional synaptic proteins involved in plasticity.

The facilitative action of lactate for the molecular interaction between GluN2B and CaMKII was validated by immunoprecipitation assays in synaptosomes and by proximity ligation assays in neurons (Figure 6). The latter observation is of particular interest as it indicates that lactate produced in the proximity of synapses through the ANLS may promote the recruitment of GluN2B containing NMDARs at the synapse, a process that has been shown to be associated with synaptic plasticity (Theodosis et al., 2008; Papouin and Oliet, 2014). It is worth keeping in mind here that the trigger for the ANLS is the release of glutamate and its subsequent reuptake by astrocytes ensheathing the synapse is the initial step of the astrocytic aerobic glycolysis that results in lactate release. Hence a sustained glutamate release as occurs during LTP induction will enhance the ANLS metabolic pathway resulting in sustained lactate release (Suzuki et al., 2011) that in turn will potentiate, through the molecular mechanisms described in this article, the post synaptic NMDAR-mediated currents with the ensuing induction in expression of genes that sustain plasticity, learning and memory (Suzuki et al., 2011; Yang et al., 2014; Margineanu et al., 2018). In this context it is worth noting that, as synaptic and cognitive loads increase, lactate becomes indispensable to sustain plasticity and cognitive performance (Dembitskaya et al., 2022).

Possibly the main message of this article is the evidence that a molecule such as lactate produced by metabolic processes known to be produced predominantly, if not exclusively, in astrocytes, is a key player in modulating synaptic transmission, an exquisitely neuronal function at the basis of learning and memory. The work presented in this article provides with molecular resolution the mechanism of this interaction further pointing at the importance of a neuron astrocyte functional unit that links energy metabolism with neurotransmission and higher brain functions.

## Author contributions

H.F. and G.H.-L. contributed equally to this work. H.F. and P.J.M. conceived the project and supervised research. H.F., G.H.-L., F.L.-C., L.M., J.G., N.C., H.M., A.O., C.B.-A., and P.J. collected, analyzed, and interpreted the data. G.H.-L. produced the figures with input from H.F. F.G. and M.D.L. oversaw and supervised PLA experiments. S.T.A. provided guidance on the mechanism of CaMKII/GluN2B interaction. H.F., G.H.-L., and P.J.M. prepared the manuscript. All authors discussed the results and commented on the manuscript.

## Data availability

All the data is available in the main text or in the supplementary materials.

---

**Figure S1.**
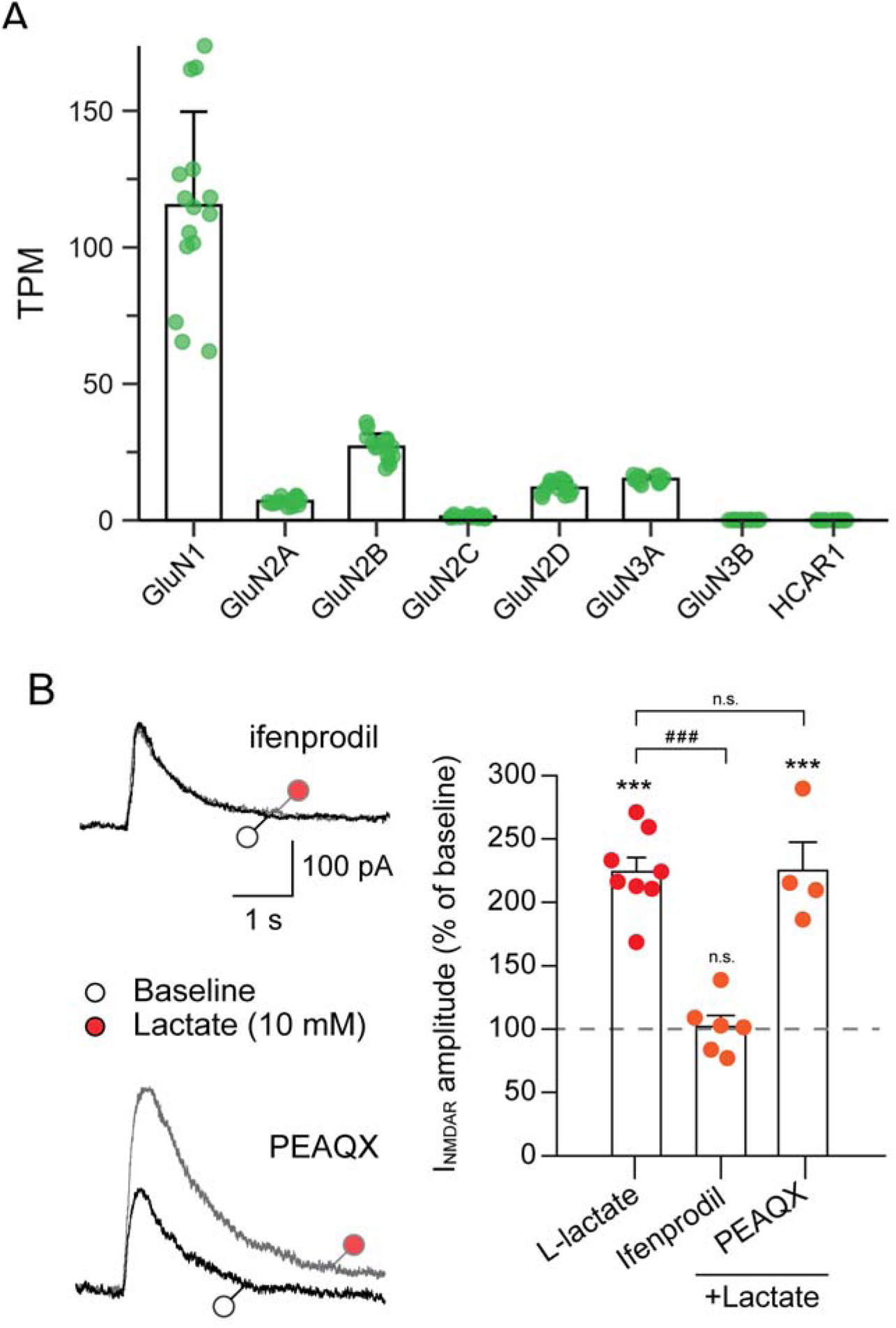
The potentiation of I_NMDAR_ by lactate is mediated by GluN2B-containing NMDARs in cultured cortical neurons. (A) Normalized mRNA expression levels of NMDAR subunits and HCAR1 in cultured cortical neurons indicated in Transcripts Per Million (TPM) using RNA-seq data (control samples) from Margineanu et al. (2018). (B, left panels) Representative traces of I_NMDAR_ responses elicited by brief applications of NMDAR co-agonists in cultured neurons recorded at a holding potential of +30 mV during the baseline (○) and 5 min after the application of drugs (●). (B, right panel) Summary bar chart showing I_NMDAR_ peak responses following bath application of 10 mM lactate in the absence or presence of ifenprodil (2 µM) or PEAQX (0.5 µM). Data are expressed as percentages of average baseline values ± SEM. Statistical significance was determined as described in Figure 1.

**Figure S2.**
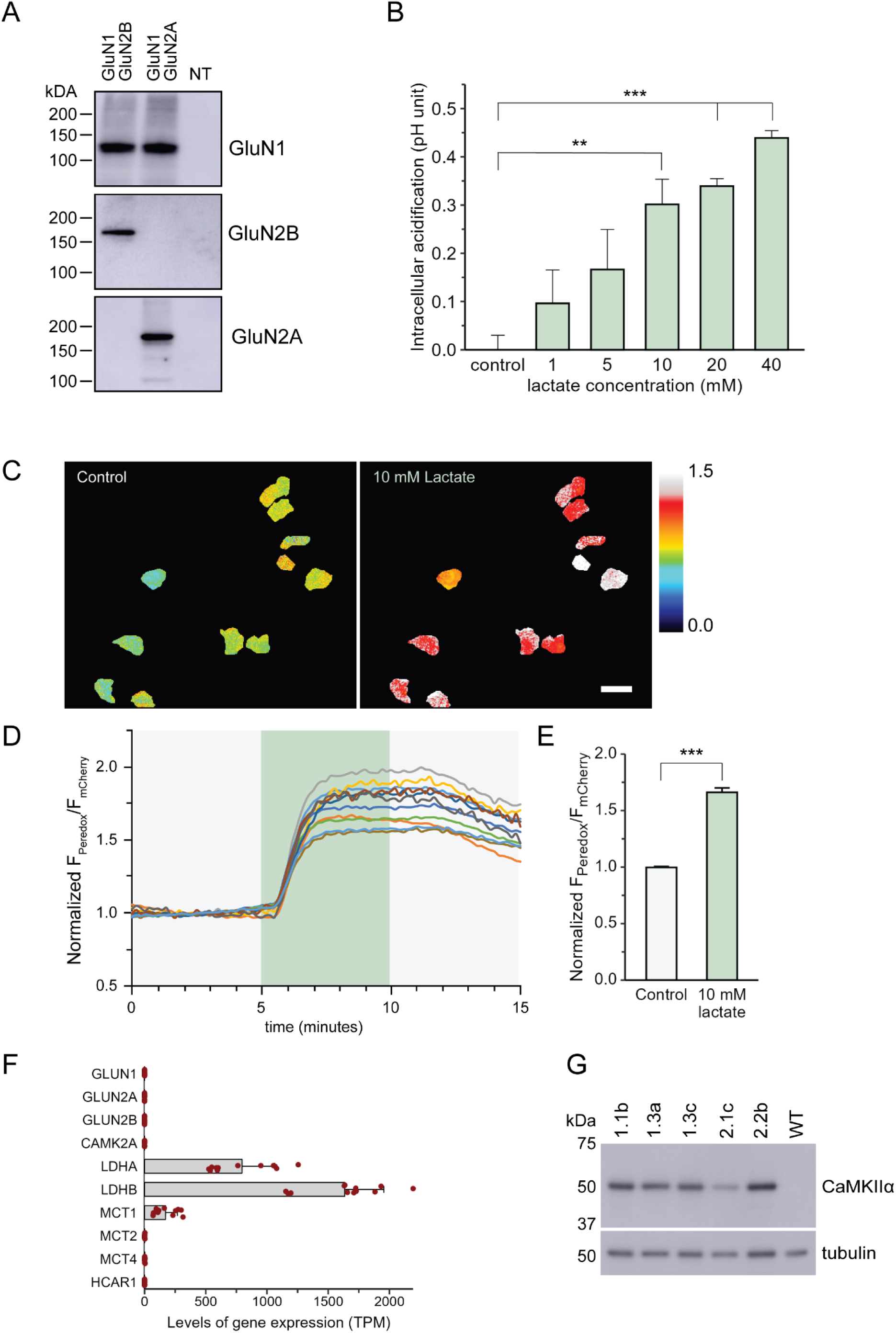
HEK cells are a suitable model to study lactate actions. (A) Western blots from untransfected (NT) HEK cells or cells transfected with GluN1, GluN2B, GluN2A subunits. (B) Bath perfusion of increasing concentrations of lactate induces an increase in intracellular acidification of HEK cells in a concentration-dependent manner. Statistical significance: ** p < 0.01, *** p < 0.001; One-Way ANOVA followed by Dunnett’s multiple comparisons test. (C) Representative images depicting the background-subtracted ratio of Peredox/mCherry fluorescence in transfected HEK cells under baseline conditions (left) and 5 minutes after the addition of 10 mM lactate (right), represented using a cold-to- warm color scale (D) Time-course of the normalized Peredox/mCherry fluorescence ratio for cells in (C), acquired every 2 seconds. (E) Quantitative summary of the averaged normalized relative F_Peredox_/F_mCherry_ fluorescence levels ± SEM, taken before and 5 min after the lactate treatment. Data are from two independent experiments. Statistical significance: *** p < 0.001, two-tailed t-test. (F) Expression levels of transcripts relevant for the lactate-induced effects in HEK cells. Note that HEK cells exhibit native expression of LDHA, LDHB, and MCT1. (G) Western blots from WT and transfected HEK cells stably expressing CaMKIIα.

**Figure S3.**
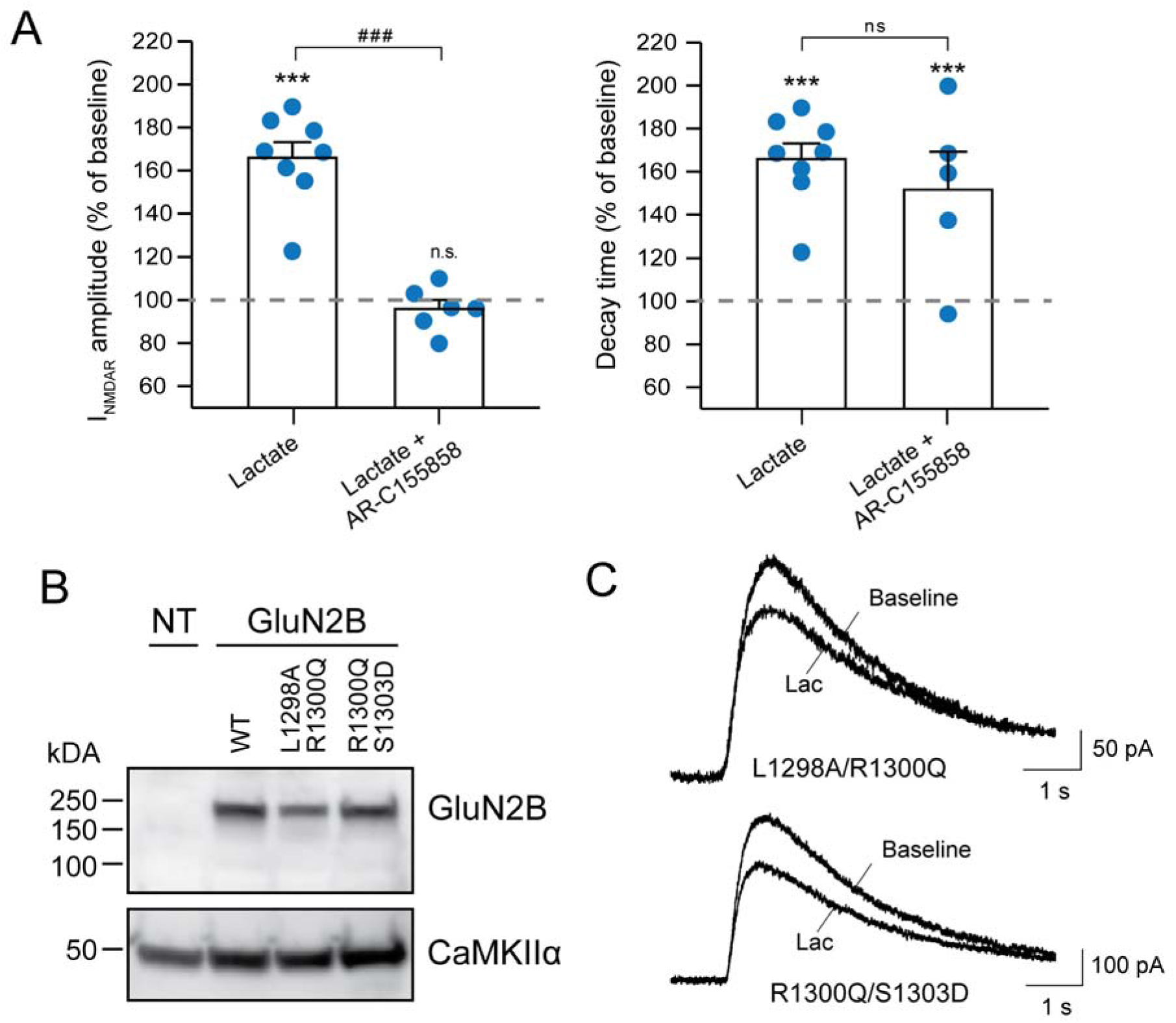
Effects of lactate and MCT blockade on NMDAR current properties and protein expression in CaMKIIα-expressing HEK cells (A) Summary bar charts of I_NMDAR_ amplitudes (left) and decay times (right) recorded at a holding potential of +30 mV in CaMKIIα-expressing HEK cells treated with lactate alone or in combination with the MCT1/2 blocker AR-C155858. Data are expressed as percentages of average baseline values ± SEM. Statistical significance was determined using paired t-tests between baseline and treatment corrected for multiple comparisons with Tukey’s method (*** p < 0.001, n.s. = non-significant), and unpaired two-tailed t-test for differences in changes across conditions (### p < 0.001, n.s. = non-significant). (B) Representative Western blot showing GluN2B and CaMKII protein expression in untransfected CaMKIIα-expressing HEK cells and these cells transfected with expressing constructs encoding mutated GluN2B subunits. (C) Representative isolated I_NMDAR_ traces recorded from CaMKIIα-expressing HEK cells transiently transfected with expressing constructs encoding the NMDAR subunits GluN1 and mutated GluN2B corresponding to the graphs shown in the figure 4D-E.

**Figure S4.**
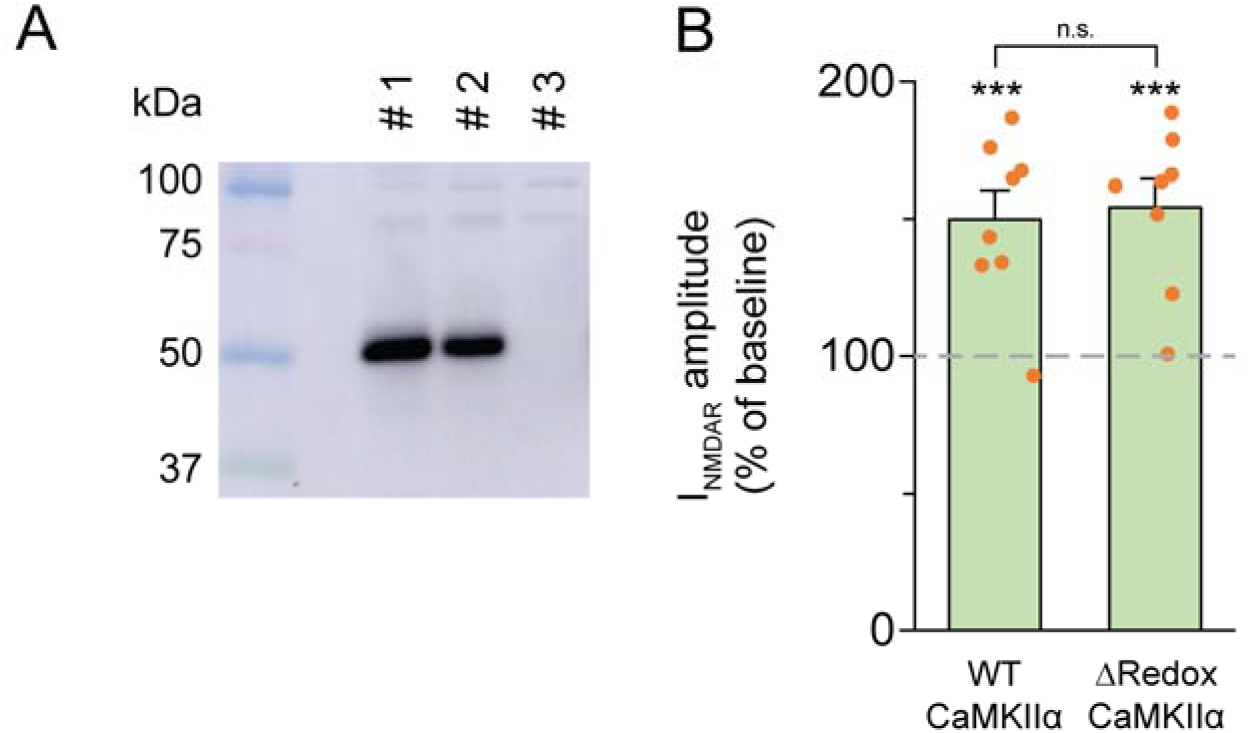
Redox-insensitive CaMKIIα does not prevent the lactate-induced NMDAR potentiation. (Left) Representative Western blot experiment showing the expression of CaMKII in two HEK cell lines (#1 and #2) stably expressing the Δredox-CamKIIα mutant compared to wild-type, non-CaMKIIα-expressing, HEK cells (#3). (Right) Bar graphs summarizing the amplitudes of I_NMDAR_ ± SEM in HEK cells expressing the WT CaMKIIα or the Δredox-CaMKIIα mutant in response to lactate treatment. Statistical significance was determined explained in Figure S3.

**Figure S5.**
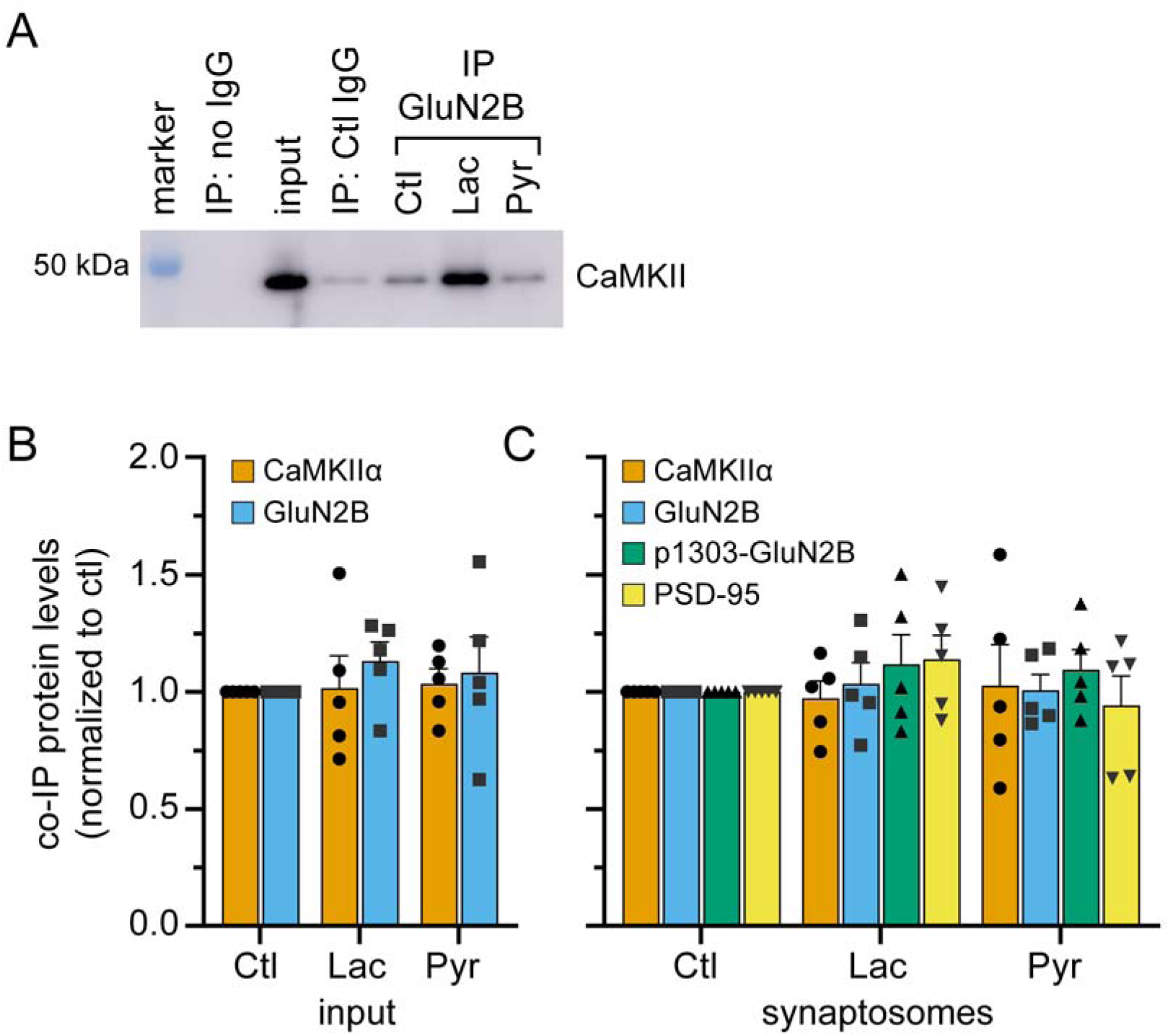
Monocarboxylates do not regulate the expression of CaMKII, GluN2B, p1303-GluN2B, and PSD-95 in the whole cell and synaptosome fractions. (A) Representative western blot showing CaMKII expression for a control co-IP experiment using no bait antibody, a control IgG or an anti-GluN2B from synaptosome fractions of 17 DIV cultured cortical neurons. The input sample corresponds to whole cell lysates from untreated neurons. Ten and one µg of proteins were loaded for the input and the synaptosome fractions, respectively. Bar graphs summarizing (B) the protein levels of CaMKII, GluN2B, p1303S-GluN2B, and PSD-95 ± SEM in control conditions or after a 20 min treatment with 10 mM lactate or pyruvate in whole cell lysate or (C) in the synaptosomal enriched fraction. Not statistically significant: Two-Way ANOVA followed by Dunnett’s multiple comparisons test.

